# Enhancing Robustness, Precision and Speed of Traction Force Microscopy with Machine Learning

**DOI:** 10.1101/2022.09.02.506331

**Authors:** Felix S. Kratz, Lars Möllerherm, Jan Kierfeld

**Affiliations:** TU Dortmund University, Germany, Department of Physics

## Abstract

Traction patterns of adherent cells provide important information on their interaction with the environment, cell migration or tissue patterns and morphogenesis. Traction force microscopy is a method aimed at revealing these traction patterns for adherent cells on engineered substrates with known constitutive elastic properties from deformation information obtained from substrate images. Conventionally, the substrate deformation information is processed by numerical algorithms of varying complexity to give the corresponding traction field via solution of an ill-posed inverse elastic problem. We explore the capabilities of a deep convolutional neural network as a computationally more efficient and robust approach to solve this inversion problem. We develop a general purpose training process based on collections of circular force patches as synthetic training data, which can be subjected to different noise levels for additional robustness. The performance and the robustness of our approach against noise is systematically characterized for synthetic data, artificial cell models and real cell images, which are subjected to different noise levels. A comparison to state-of-the-art Bayesian Fourier transform traction cytometry reveals the precision, robustness, and speed improvements achieved by our approach, leading to an acceleration of traction force microscopy methods in practical applications.

SIGNIFICANCE
Traction force microscopy is an important biophysical technique to gain quantitative information about forces exerted by adherent cells. It relies on solving an inverse problem to obtain cellular traction forces from image-based displacement information. We present a deep convolutional neural network as a computationally more efficient and robust approach to solve this ill-posed inversion problem. We characterize the performance and the robustness of our approach against noise systematically for synthetic data, artificial cell models and real cell images, which are subjected to different noise levels and compare performance and robustness to state-of-the-art Bayesian Fourier transform traction cytometry. We demonstrate that machine learning can enhance robustness, precision and speed in traction force microscopy.

## INTRODUCTION

Many cellular processes are intrinsically connected to mechanical interactions of the cell with its surroundings. Mechanical surface forces control the shape of single cells or groups of cells in tissue patterns and morphogenesis (1). Forces alter cell behavior via mechanotransduction (2) and affect cell migration and adhesion. Gaining access to the forces (or tractions, i.e., forces per area) exerted by the cell during critical processes like migration or proliferation can give insight into biophysical processes underlying force-generation and aid the development of novel medication and treatment, e.g. by identifying changes of cellular forces in diseased states. Altered cell behavior is present for diseases (3) like atherosclerosis (4), deafness (5), or tumor metastasis (6).

*T*raction *F*orce *M*icroscopy (TFM) is a modern method to measure tractions exerted by an adherent cell by deducing them from the cell-induced deformations of an engineered external substrate of known elastic properties (7–9). Beyond adherent cells it has applications to a broader range of biological and physical systems where interfacial forces are of interest (10). TFM thus constitutes a classic inverse problem in elasticity, where tractions or forces are calculated from displacement for given material properties. This inverse problem turns out to be ill-posed, i.e., noise or slight changes in displacement input data induce large deviations in traction output data because of singular components of the elastic Green’s tensor. This technical problem has been addressed by different regularization schemes that have been developed over the last two decades (11–15). Recent studies show that *M*achine *L*earning (ML) can be an elegant alternative to numerical schemes when the inverse problem to a bounded problem is ill-posed in the context of elasticity or rheology. This has been explored for other rheological inverse problem classes such as in pendant drop tensiometry (16). ML-aided traction force determination can thus provide an elegant way to improve TFM as a method, as recent studies have already begun to show (17, 18). A systematic investigation of ML-aided TFM with respect to an optimal general purpose training set that allows the machine to predict tractions accurately across many experimental situations as well as a systematic investigation of accuracy and of robustness with respect to noise, which is present in any experimental realization, are still lacking.

The first implementation of TFM was achieved by Harris and coworkers in the early 1980s, where thin silicone films are wrinkled by compressive surface stresses, inflicted by the traction field of the cell (19). Due to the inherent non-linearity of wrinkling and the connected difficulties solving the inverse elastic problem, this method has been superseded by linear elastic hydrogel marker based TFM introduced by Dembo et al. (20). Due to the simplicity of the hydrogel marker based approach, it is the most commonly used and most evolved method. Alternative techniques and extensions include micro-needle deformations (21), force microscopy with molecular tension probes (22), and 3D techniques (23). Wrinkling based TFM has recently been re-explored with generative adversarial neural networks with promising results (18).

In this paper, we focus on the hydrogel marker based technique and train a deep *C*onvolutional *N*eural *N*etwork (CNN), which has the capabilities to solve the inverse elastic problem reliably, giving fast and robust access to the traction pattern exerted by the cell onto a substrate. Specifically, we do this by numerically solving the elastic forward problem, where we prescribe generic traction fields and solve the governing elastic equations to generate an associated displacement field. The “synthetic” displacement field generated this way is used as a training input for our NN, while we use the prescribed traction field as the labels for our training set. This way the network learns the mapping between displacement and traction fields and is able to generate traction fields for displacement fields never seen before, while still respecting the relevant governing elastic equations. Complete knowledge of the prescribed tractions for the synthetic training data enables a training process that directly minimizes deviations in the predicted tractions. This contrasts conventional TFM techniques which determine traction forces indirectly by minimizing deviations in the resulting displacement field. We use traction force distributions generated from collections of circular force patches as training data, which seems a natural general choice to allow the NN to predict generic force distributions in cell adhesion but should also cover other future applications. We show that the proper, “physics-informed” choice of training data and inclusion of artificial noise is a similarly important step in the ML solution of the inverse problem as the proper choice of regularization in conventional TFM techniques, in order to achieve the best compromise between accuracy and robustness.

## MATERIALS AND METHODS

### Hydrogel Marker Based TFM

The hydrogel marker approach to TFM can be described as follows. First, a cross-linked gel substrate, often Polydimethylsiloxane (PDSM) or Polyacrylamide substrates (PAA) (24), is cultivated. The cross-linked gel can be classified as an elastic substrate with long linkage lifetimes compared to the imaging process (25).

Second, the substrate is coated with proteins prevalent in the extracellular matrix like collagen type I, gelatin, laminin, or fibronectin, allowing the cell to adhere to the substrate. Fluorescent marker beads embedded in the gel substrate aid the determination of cell-induced substrate deformations. The reference and stressed positions of the marker beads can be determined via various microscopy techniques, ranging from confocal to optical microscopy (19).

Third, to infer the displacement field from the marker bead positions, a particle tracking velocimetry (PTV) algorithm, a particle image velocimetry (PIV) algorithm or a CNN particle tracker (26) is used, which calculates the discrete displacement field.

The information about the displacement field, combined with the predetermined constitutive properties of the hydrogel substrate gives access to the traction field of the cell via the inverse solution of the elastic deformation problem. For homogeneous, isotropic, and linear elastic solids the displacement field 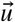 satisfies the equations of equilibrium in the bulk (27)

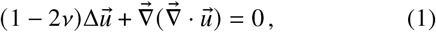

while the force balance at the surface is modified to account for external tractions 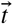 (forces per area applied to the surface)

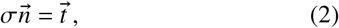

where 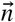 is the surface normal vector and σ the stress tensor.

The TFM gel substrate can be considered sufficiently thick to be modelled as an elastic half-space (*z* > 0), bounded by the *x*-*y*-plane, at which traction forces 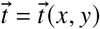 are applied. The displacements are a solution of the boundary problem eqns. (1), (2), which is given by the spatial convolution of the external traction field 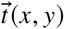 with the Green’s tensor **G** over the boundary of the surface *S* (27):

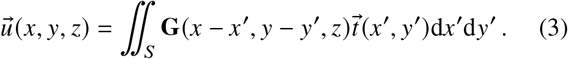

Up to this point both tractions and displacement are threedimensional vectors. The full three-dimensional Green’s tensor **G** (*x* − *x*^′^, *y* − *y*^′^, *z*) is given in the Supporting Material.

In TFM, it can be assumed that adherent cells exert *inplane* surface tractions (*t*_*z*_ = 0), and we are interested in *in-plane* displacement 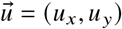 only because out-of-plane *z*-displacements are hard to quantify by microscopy. Moreover, out-of-plane displacements are small for incompressible materials (see Supporting Material). These assumptions make the problem eqn. (3) quasi-two-dimensional in the plane *z* = 0, such that the Green’s tensor is given by the 2×2 matrix (20)

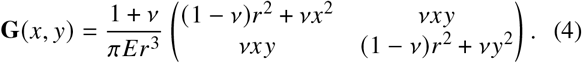

It solves the elastic boundary problem for in-plane tractions and displacements if the tractions vanish at infinity.

TFM is essentially a technique to provide a numerical solution for the inverse elastic problem posed by asking to recover the traction field 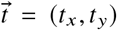 from eqn. (3) via a deconvolution of the right hand side surface integral. This can be done in real space (11, 12, 20) or in Fourier space (28).

Employing the convolution theorem for the Fourier-transform ℱ 𝒯 of a convolutional integral, the deconvolution problem encountered in eqn. (3) can equivalently be stated as performing two Fourier-transforms and one inverse Fourier-transform

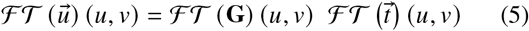

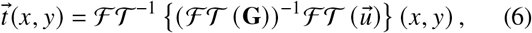

which is named *F*ourier-*T*ransform-*T*raction-*F*orce-*C*ytometry (FTTC) (28).

Common iterative techniques used for numerical deconvolution can become unstable when subjected to noisy data, which is why conventional approaches to the ill-posed inverse elastic problem rely on regularization techniques (e.g. Tikhonov(L2)- or Lasso(L1)-regularization) coupled with iterative minimization schemes (11–14, 29). This applies both to real space and Fourier space methods. These methods minimize deviations in the resulting displacement field subject to suitable regularization constraints on the traction forces. Regularization improves stability while accuracy might suffer. The optimal choice of regularization parameters is important but subjective. In *B*ayesian *F*ourier *T*ransform *T*raction *C*ytometry (BFTTC) the regularization parameters need not be picked manually and heuristically, but they are inferred from probability theory, making an easy to use and objective FTTC method (15, 30).

The shortcomings of most conventional approaches are systematic under-predictions and edge smoothing of the constructed traction field, caused by the regularization (31), as well as elevated computational effort, inflicted by the computationally demanding iterative deconvolution techniques and transformations at play.

A recent trend in many fields, including the natural sciences, has shown the capabilities of ML-based approaches in such ill-posed and ill-conditioned scenarios (32, 33), often outperforming complex algorithms by orders of magnitude in computing time and precision, and thus allowing for new and more accessible workflows with reduced computational overhead. ML based approaches to TFM (17) and wrinkle force microscopy (18) have recently been discussed and find that deep CNNs can perform the deconvolution of eqn. (3), by learning the mapping from strain-space to surface traction-space in training. The existing NN approaches show a promising proof-of-concept which we want to extend further in the present article by systematic studies of accuracy and robustness to noise. While regularization is particularly important in conventional TFM approaches for accuracy and stability, accuracy and robustness to noise of deep CNNs crucially depend on the choice of training data.

“Physics-informed” ML methods have also been applied to directly solve general partial differential equations with boundary conditions, such as eqns. (1) and (2) that are underlying TFM (34–36). In TFM we can solve the elastic problem analytically up to the point that the Green’s tensor eqn. (4) is exactly known but proper inversion is difficult. We want to solve this inversion problem by deep CNNs with a “physics-informed” choice of training data and learning metric.

### Machine Learning the Inverse Problem

If we want to teach a machine to solve an inverse problem for us, counter-intuitively, we do not need to know how to solve the inverse problem itself. We only need to know how to solve the corresponding forward problem i.e., we only need to know how to precisely formulate the learning task for the network and provide sample data that characterize the problem well enough. In training, the machine detects correlations in the data and uses it to solve for arbitrary non-linear mappings from input space to output space. This is one of the groundbreaking traits deep learning offers and which, in combination with hardware acceleration, allows to solve problems not feasible or traceable before (16, 32, 33).

Thus, we are interested in formulating the learning task in the most precise way. A first step to this is a solid understanding of the forward problem and all involved quantities including the proper, “physics-informed” choice of training data. Since we generate our training data numerically, the second step is a robust numerical framework, which outputs physically correct data. The third step is to find a sensible data representation, which contains all the relevant information. The final step is to find a network architecture with enough capacity for the problem, to train it and to evaluate its performance and robustness. We will address these points in the following and finally compare our network with state-of-the-art approaches.

#### Understanding the forward problem for traction patches

The forward problem we are trying to solve involves cell tractions on elastic substrates. Thus, essential to the performance of our NN is the accurate interpretation of cell-characteristic deformations of the substrate. Cells generate forces exerted onto the substrate via focal adhesion complexes with sizes in the *μ*m-range and tractions in the range nN *μ*m^−2^ = kPa (37). Forces are generated by tensing acto-myosin stress-fibers that attach to the focal adhesions and, therefore, have a well-defined direction over a focal adhesion complex. Therefore, typical cellular traction patterns consist of localized *patches*, which can comprise single or several focal adhesion complexes and are anchored to the substrate at positions 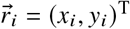. Within these patches tractions have a well-defined in-plane angle *γ*_*i*_ with the *x*-axis resulting in a traction pattern

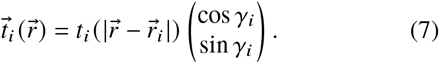

These tractions are applied to circular patches of variable radius *R*_*i*_ at the anchored nodes (see Fig. 1), such that

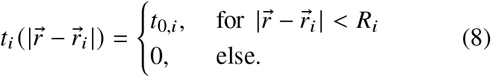

**Figure 1:**
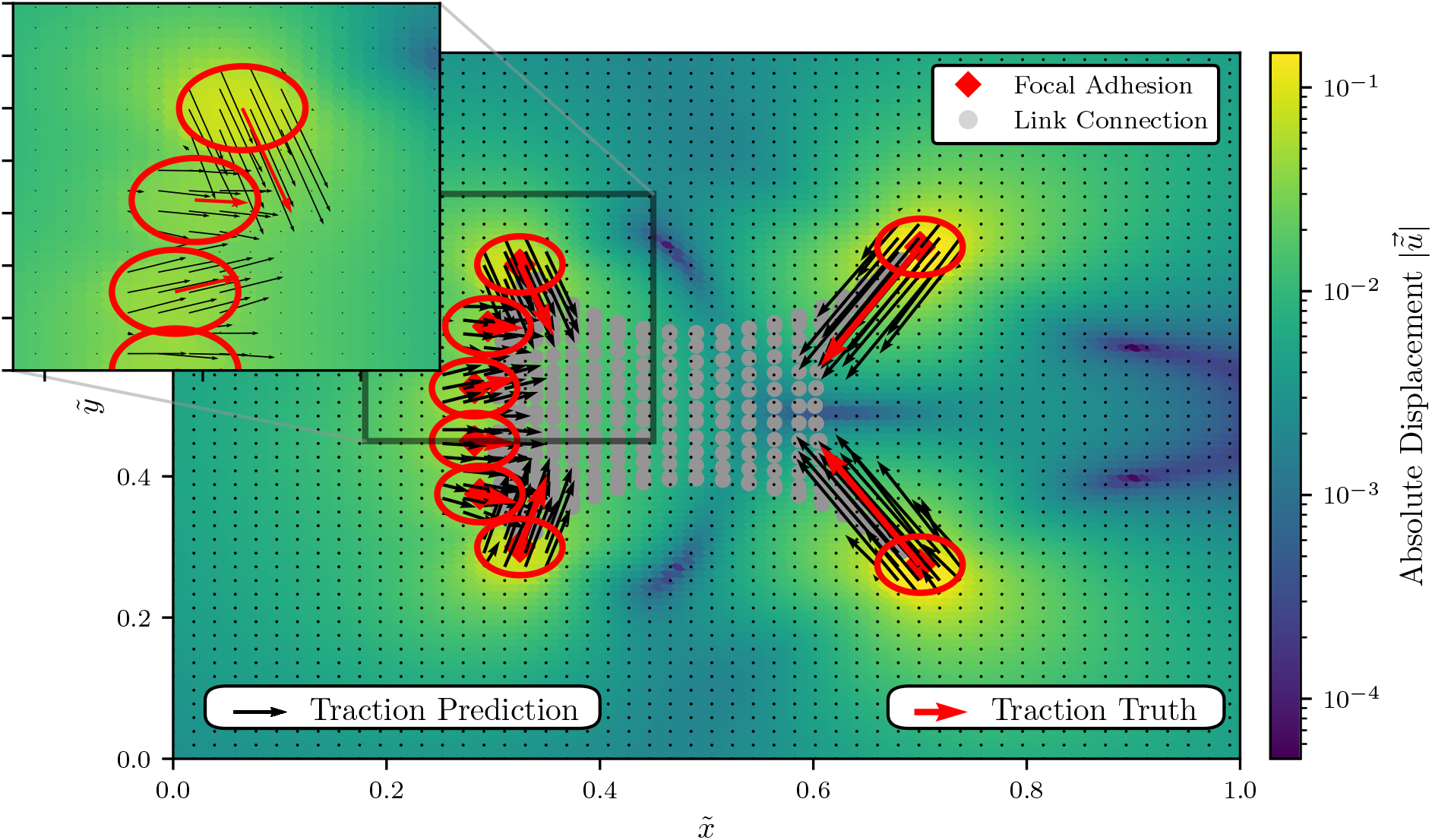
A model cell with circular focal adhesion points. The model cell perturbs the elastic substrate it is resting on by generating tractions (red arrows) at the focal adhesion spots (red circles), resulting in the color coded displacement field; tractions (red arrows) are generated based on a contractile network model (see text). Red arrows are the “true” average tractions generated by the cell model over the red circles, while the black arrows indicate the local tractions that the NN_low_ network predicts at the discrete grid spots.

We model typical traction patterns as a linear superposition of traction patches eqn. (7) localized at different anchoring points. Within linear elasticity, the resulting displacement pattern is also a linear superposition of all the displacement patterns 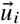 caused by all traction patches *i*.

For a single traction patch, we solve the forward elastic problem by exploiting the convolution theorem

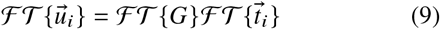

where the Fourier transform of the Green’s kernel in polar coordinates *ρ* and *ϕ* is known (28),

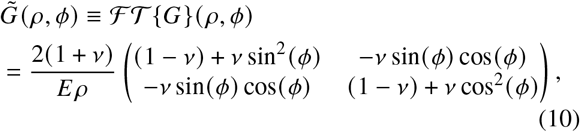

and the Fourier transform of the traction spot is given by the Hankel transform

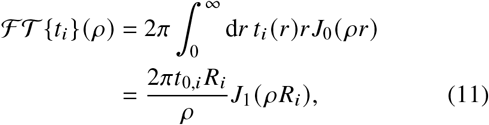

where *J*_*n*_ are the Bessel functions of the first kind.

The Fourier transformed displacement field 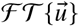 is now accessible and can be converted back to the displacement field by performing the inverse Fourier transform 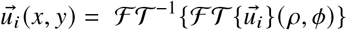. The inverse Fourier transform can be performed analytically in polar coordinates centered around the corresponding anchored node with a scaled radial component 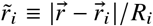 and an angle *θ* with the *x*-axis,

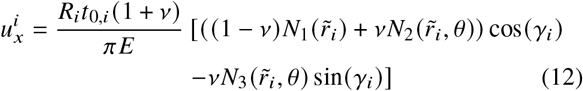

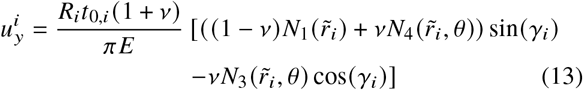

where *N*_1,2,3,4_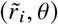 are specific functions, that describe the geometric dependence of the displacement field, and are obtained by explicitly solving the occurring inverse Fourier transforms in the Appendix (see eqns. (19), (20), (21), (22)). Strictly speaking, this analytical solution of the forward elastic problem for a single traction patch anchored at *R*_*i*_ is valid on an infinite substrate. We will neglect finite size effects in the following, and use this analytical solution also on finite substrates. The solution for many traction patches anchored at different points is obtained by linear superposition.

#### Numerically solving the forward problem

We consider a square substrate of size *L*×*L*, in which displacements are analyzed (the total substrate size can be larger). Typical sizes are in the range *L* ∼ 10 − 100*μ*m. We use the size *L* to non-dimensionalize all length scales: 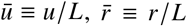 and 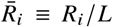, such that the substrate in which displacements are observed always has unit size. Typical focal adhesion patch sizes *R*_*i*_ are in the range of several *μ*m (15, 37); in dimensionless units, we take 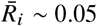 as typical value. The above dimensionless coordinate 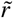 remains unchanged by nondimensionalization. Furthermore, we use the elastic constant *E* as traction scale: 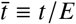. Typical hydrogel substrate elastic moduli of *E* ∼ 10kPa (15) and tractions in the range up to 5kPa (37) imply typical dimensionless tractions up to 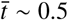.

We create a *N*×*N* square grid, on which we discretize the solution of eqns. (12) and eqn. (13) for a supplied traction patch 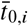 with direction *γ*_*i*_ and use superposition of the individual patch solutions for all anchored nodes, such that we get the full displacement field for a number of *n* circular traction patches of variable radius *R*_1,…,*n*_.

We discretize both displacement and traction fields on the same *N* × *N* square grid. While generating the displacements in eqns. (12) and (13) on a discrete grid is simple, we note that the discretization of the traction field needs to be performed with great care. A naive approach for the discretization of the circular traction patches onto a square pixel grid with indices *i, j* ∈ {1, …, *N*} would be the direct discretization of eqn. (7), i.e., to check whether any square segment center point 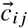 is contained in the circular traction patch of radius *R*_*t*_ and center point 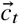. If the center point is contained, the grid segment *i, j* is assigned the traction 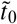 of the circular patch. This naive discretization suffers from a critical artifact: it is not force conserving, i.e., does not conserve the total traction force exerted by the patch, which is given by the area-integrated tractions. This violates the fundamental physical requirement of force balance.

In the Supporting Material, we discuss improvements and present an exactly force conserving traction discretization procedure by calculating the *exact* overlap area *A*_ov_ of each square grid segment (with side lengths *a* = *L* / *N*) and the circular traction spots. Then we assign a corresponding fraction 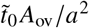 of the traction to each square grid segment. In the Supporting Material, we also quantify the accuracy gain by the force conserving traction discretization method by computing the errors in the displacement field of a large circular patch that is discretized.

We want to emphasize the relevance of these findings to our approach: As the NN will be trained with the discretized traction fields and we are ultimately interested in an accurate *discretized* traction prediction by our machine, we are forced to deliver as accurate discrete traction field representations as “truths” for training as possible.

#### Generating arbitrary traction fields via superposition

Because of the underlying linearity of the elastic problem at hand we are able to construct displacements for *arbitrary* traction patterns via superpositions of the circular traction patch solutions. While this might seem obvious, it has far reaching implications for our approach and implies that a solver with the ability to reconstruct traction fields constructed from circular traction patches will also be able to reconstruct *arbitrary* traction fields if the solver preserves the linearity of the elastic problem.

Because we will present our NNs with an arbitrary superposition of circular traction spots, discretized to a finite grid, it is trained to exploit the linearity of the problem explicitly, and we thus expect the networks to be able to solve the more general problem of predicting an arbitrary superposition of traction patches. In a sense, generating superpositions of the analytical circular traction patch solutions is an optimization we employ to reduce the computational effort for generating displacement fields for training, while retaining the relevant properties of the problem, as we will show.

Another implication of this observation is that we are able to check the predicted discretized traction fields for consistency with a supplied displacement field by constructing a superposition of displacement fields for circular traction spots with radius 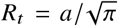 for each grid point, where is the distance between grid points. The choice 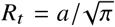 assures conservation of the total traction force. We implement this method along with our solver to generate the displacement fields from arbitrary superpositions of circular traction patches.

#### Architecture of the deep convolutional neural network

We choose to employ a *Unet* structure (38) consisting of an *input* encoder, which extracts and compresses the relevant information from the high-dimensional *input* displacement field into a lower-dimensional representation. From the lower-dimensional and compressed displacement information we inflate the dimensionality again with a decoder, such that we finally receive the representation of a traction field in the *output* of the network, as shown in Fig. 2. The motivation for this choice is the conceptual similarity of image procession tasks such as segmentation, which involve local classification of an image, to the assignment of local “traction labels” to each grid point of the “displacement image”. Furthermore, the elastic problem has long range interactions, where a localized traction spot causes large scale displacements. The layered structure of a Unet is well suited to handle this problem, as the high-dimensional layers process short scale information and the increasingly lower-dimensional layers will be able to handle longer range correlations. Finally, through the process of compressing and reinflating dimensionality we might loose spatial precision and, thus, use the skip connections to provide the upsampling layers with additional spatial information. Additionally, skip connections have been demonstrated to improve generalization potential and stability when used in combination with batch normalization (which is also used in our networks) (39).

**Figure 2:**
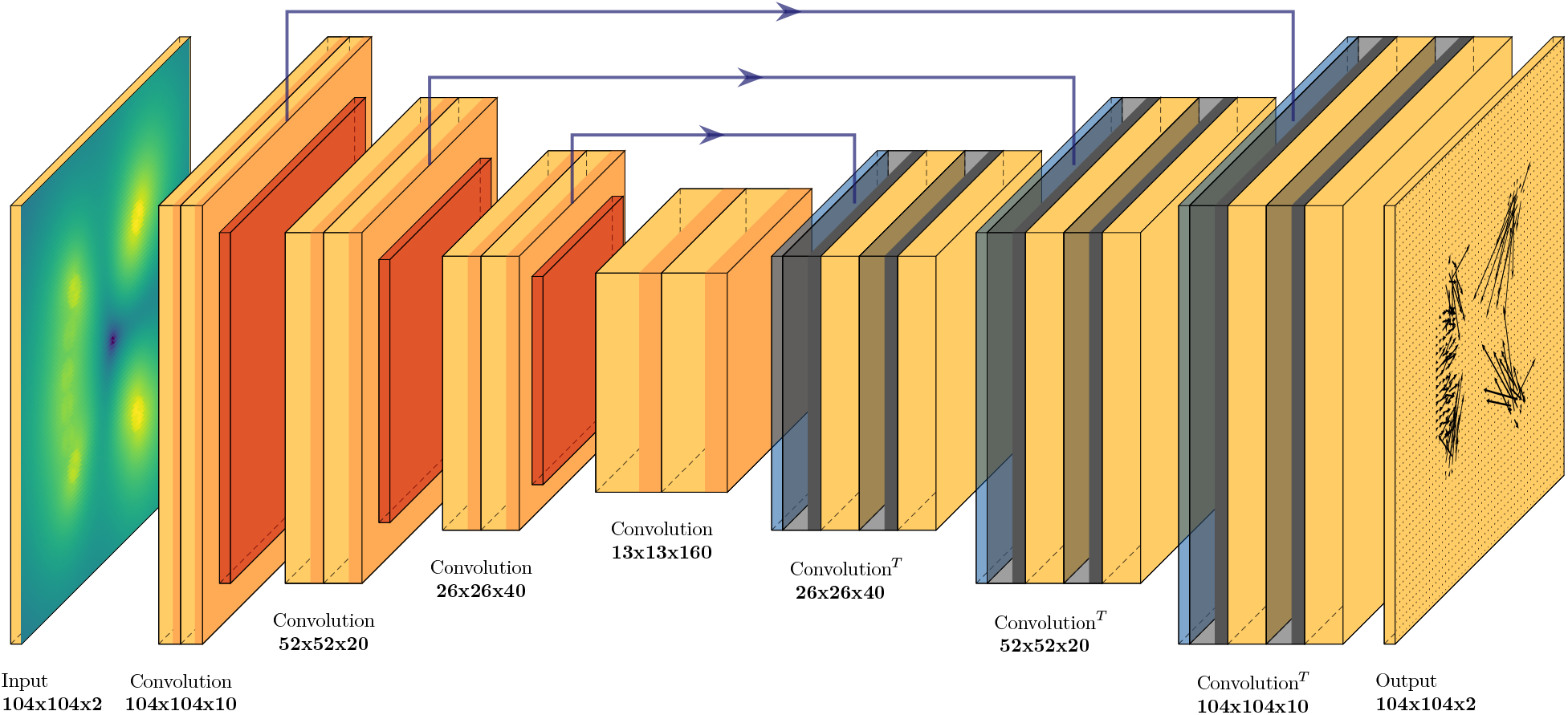
The network we employ is a *Unet* Convolutional Neural Network with a discretized displacement field as an input and a discretized traction field as an output. The mapping from input to output is learned in training by adapting the parameters of the convolutional and transposed convolutional layers of the network. Eventually the network will be able to reconstruct the traction field for displacement fields never seen before. We do not enforce a strict bottleneck, rather we allow for skip connections from the encoding process to the decoding process (blue arrows). The skip connections thus offer a way for the network to manipulate the decoding process with selected information gathered during encoding, increasing the capacity of the network.

Our network (as shown in Fig. 2) is a fully convolutional NN, where the encoding part is a stack of convolutional blocks, and max-pooling layers, while the symmetric decoding part consists of transposed convolutional layers, skip connections and convolutional blocks. Each convolutional block consist of two convolutional layers, with one *Dropout* layer and *LeakyReLU* activation functions, which introduce non-linearity. Specially, the last layer uses a linear activation function, ensuring that the output maps to the domain of a traction field. The encoding part uses size 3 kernels and size 1 strides, while the decoding part uses size 4 kernels and size 2 strides to avoid checkerboard effects that would otherwise negatively impact performance.

#### Training data sampling and the training process

To train our NN we choose a 104×104-grid (*N* = 104) which holds the discrete representation of the dimensionless displacement and traction fields. We will later show that our networks are still able to work on arbitrary grid sizes (with proper scaling of the input), since they are fully convolutional. The training data is generated by numerically solving the explicit forward problem in dimensionless form as outlined in the section Numerically solving the forward problem.

The traction distribution 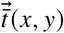 is generated by a random number (uniformly sampled in [10, 50]) of traction spots 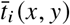, where the radius 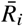 is drawn uniformly in the range [0.01, 0.05] with a random center point 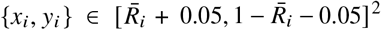. The traction magnitude 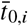 is uniformly distributed in the range [0, 0.5] and the polar angle *γ*_*i*_ is uniformly distributed in [0, 2*π*].

While traction values and patch sizes are typical for adherent cells, our training data is more general in the sense that other important characteristics of cellular force patterns, such as the occurrence of force dipoles at the end of stress fibers, are *not* contained in our training data. This makes our approach more general compared to Ref. (17), where training was performed on traction patterns typical for migrating cells. In combination with non-dimensionalization, this will allow us to easily adapt the training process to other applications of TFM in interfacial physics (10) in future applications. Below, we will demonstrate the ability of the CNN to specialize from our general patch-based training set to artificial and real cell data. As a convenient model to generate realistic cell traction data artificially we use the contractile network model of Ref. (40).

We expect a NN trained with noisy data to also perform better when confronted with noisy data. To test this hypothesis, we add different levels of background noise to the displacement field *ū* in our training data. In order to evaluate the effects on robustness we train two types of NN:

- A network NN_low_ is trained with a low level of background noise: To each dimensionless training displacement field value 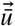 a spatially uncorrelated Gaussian noise with a variance 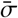 that is 0.5% of the average variance of the dimensionless displacement field over all training samples,

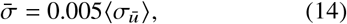

where ⟨… ⟩ is an average over all training samples.
- A network NN_high_ is trained with a high level of background noise that is 5% of the average variance of the dimensionless displacement field over all training samples,

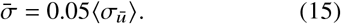

We want to emphasize that we use *uniform* Gaussian noise for the training. The assumption of uniform Gaussian noise is used as the central assumption in BFTTC to evaluate the likelihood. The training data can easily be adapted to contain different types of noise if there is a concrete experimental motivation to do so.

As a loss or performance metric we use the mean-square error (MSE) between the output guess for tractions and the corresponding labels of the input traction data averaged over *M* training batches,

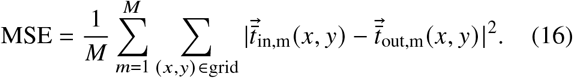

We train in batches of 50 samples per batch by backpropagation using the Adadelta algorithm. The number of training steps per epoch consists of 900 training batches or 45000 samples.

A traction-based objective function comparing the residual of the force balance that generates the displacements is the proper “physics-informed” error metric, since we are interested in correct traction forces in TFM. Training for correct traction forces is enabled by using synthetic training data based on traction patches, where we know the true tractions. Alternatively, one could use the residual between the input displacement field and a displacement field generated from the predicted traction field as a training metric, but this approach has an obvious problem: to do the backpropagation during training we would have to compute *all* predictions of the network for the displacement field in each step of the training, which slows down training several *orders of magnitude* (in a first approximation by a factor of *N*^2^). Implicitly, conventional TFM techniques such as the BFTTC algorithm follow this strategy as they minimize deviations in the resulting displacement field (subject to additional regularization) (15). Therefore, we expect networks trained according to this strategy to perform qualitatively similar to the BFTTC algorithm. We will investigate in detail the resulting differences in accuracy of the traction and displacement predictions in the section Displacement versus traction error.

During training, we evaluate the loss MSE eqn. (16) for the training data and a validation MSE for unknown displacement data of the same type. The validation and training errors in Fig. 3 show constant learning and generalization of the model without over-fitting. We note that a valuation loss lower than the training loss is common when using dropout layers, which are active in training but inactive during inference. In total, the training is performed for 5000 epochs, which we chose as an arbitrary training limit to truncate the power law tail seen in training, took ∼ 100 h on an *NVIDIA QUADRO RTX 8000* GPU, with the main learning advancements occurring in the first 5 h. Each epoch consists of 50000 randomly chosen traction patch distributions of which 45000 samples are used in training and 5000 are used for validation.

**Figure 3:**
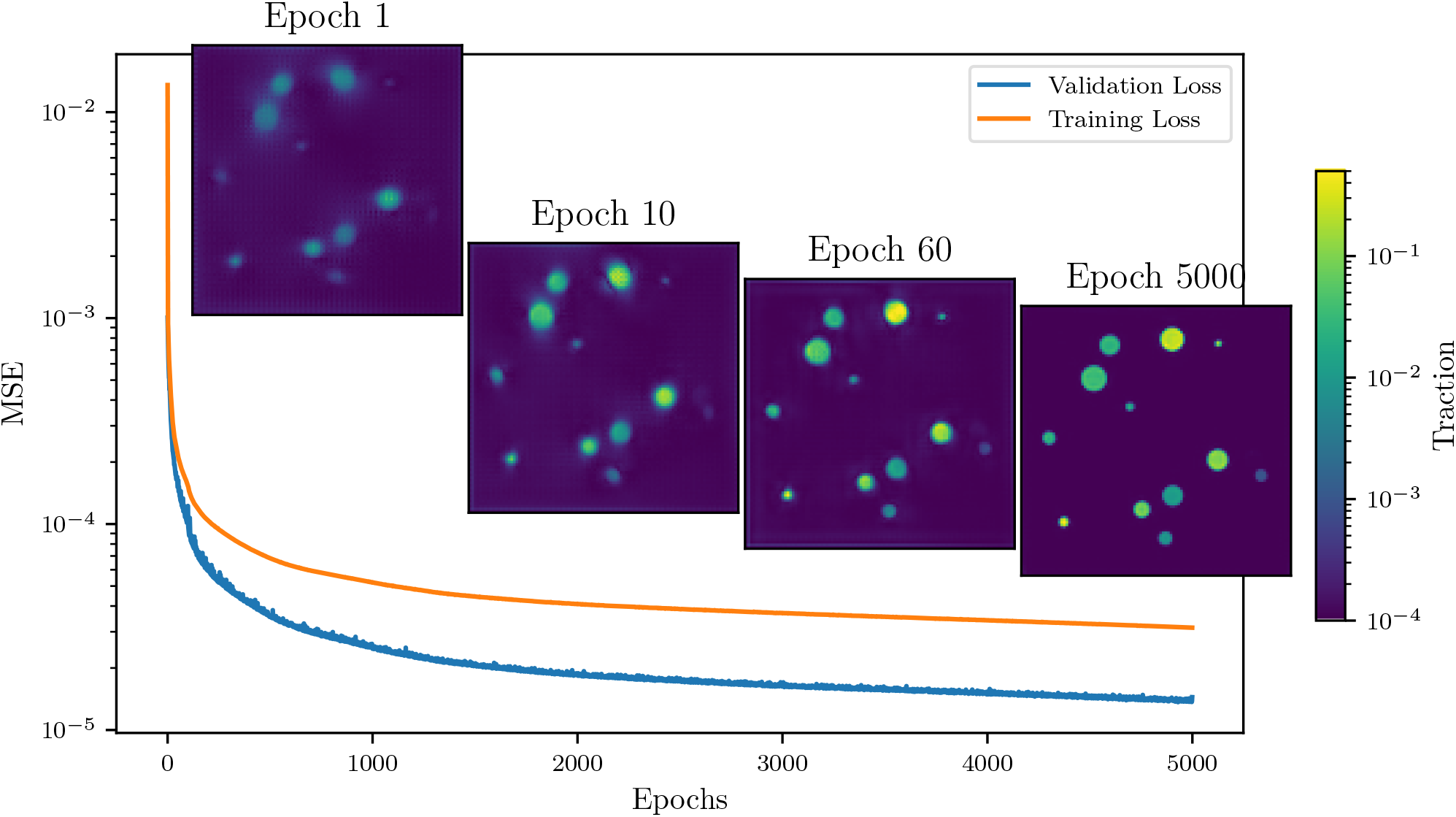
The time evolution of training and validation MSE during training of the neural network NN_low_. Already after the first epoch a rough traction reconstruction is achieved. After 10 Epochs the reconstruction gains significantly in terms of visual sharpness, which is further increased in the following epochs. We stop the training process after 5000 epochs, because longer training yields diminishing returns.

#### Sampling the hyper-parameter space of networks

We sample the hyper-parameter space to detect which network traits are important for its performance, i.e., we train a number of different networks on the same data for the same duration and compare their learning progress. The findings of this sampling are contained in Fig. 4, where the baseline (solid blue, label “Single Conv.”) is a network with a single convolutional layer per block, skip connections, a dropout of 10% and a batch size of 132, using the *Adadelta* optimizer. For this figure we use data generated with the “naive” traction patch discretization and switch to the force conserving method for one of the trials.

**Figure 4:**
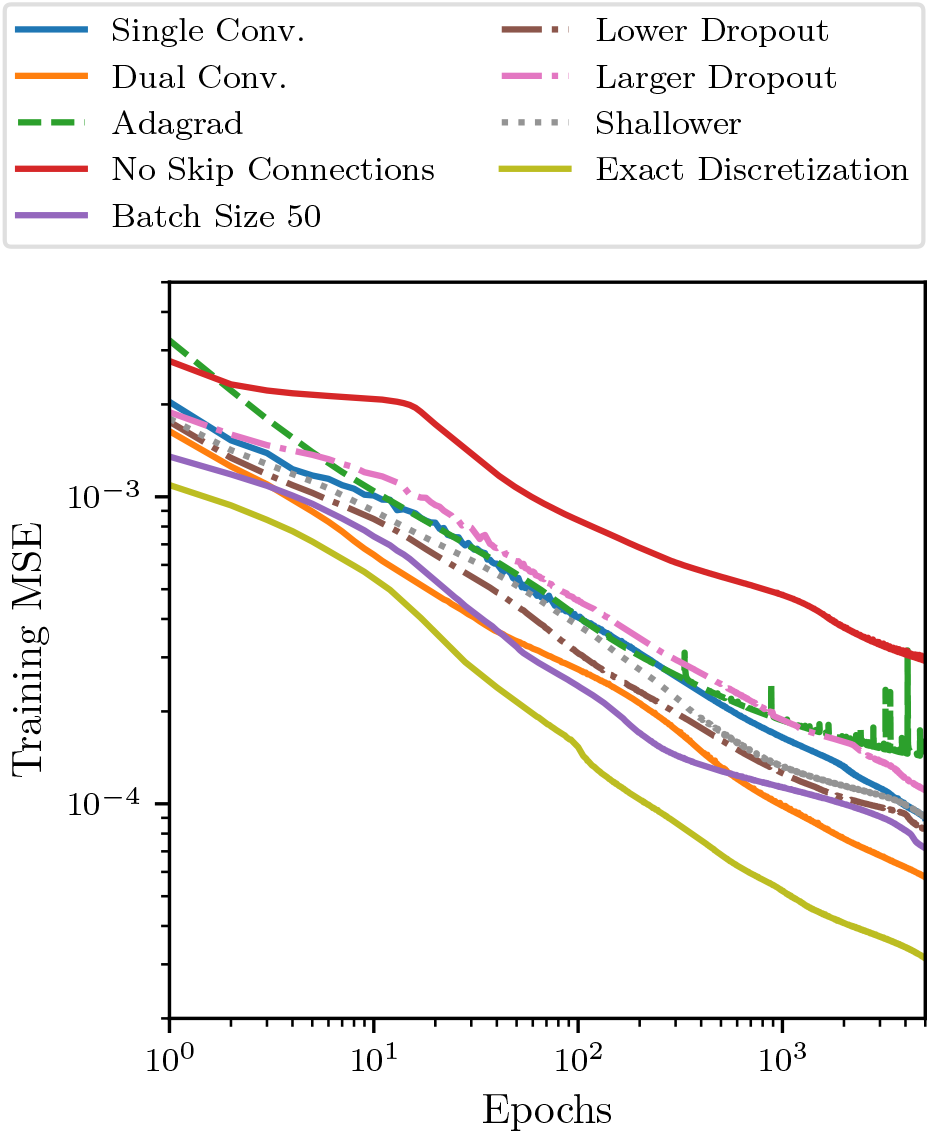
The learning processes of different networks where we sweep the hyper-parameter space by changing one property at a time to search for well performing networks (see text). Generally, training follows a power law and we cut off training at 5000 epochs.

In Fig. 4 we vary only one parameter at a time, i.e., we disable the skip connections, vary the batch size, change the optimizer, change the dropout rate, change the number of convolutional layers in a block or change the traction discretization method, this allows us to get an understanding of the hyper-parameter space and its implications on predictive performance:

1. The base line configuration “Single Conv.” (solid blue) with a single convolutional layer and trained with the Adadelta optimizer performs better compared to training with the Adagrad optimizer (“Adagrad”, dashed green).
2. The training performance is improved when using two convolutional layers per block (“Dual Conv.”, solid orange), but, because of the increased complexity, training and inference is computationally significantly more expensive.
3. We observe that the network without skip connections (“No Skip Connections”, solid red) performs significantly worse than all other networks.
4. We are able to improve learning by using a lower batch size of 50 (“Batch Size 50”, solid purple).
5. Changing dropout affects training as one would expect – a larger dropout decreases and a lower dropout increases training precision (“Larger/Lower Dropout”, dashed dotted magenta/brown).
6. When employing a shallower network obtained by removing one encoder and one decoder block (“Shallower”, dotted grey) the learning is faster initially, but seems to plateau earlier before improving again.
7. Finally, when using the exact force conserving discretization for the traction grid (“Exact Discretization”, solid yellow) we are able to drastically improve training performance, supporting our above claim that conserving the force balance exactly is of great importance.

For all network variants, the training progress shown in Figs. 3 and 4 follows a power law MSE ∝ epochs^−*α*^ with exponents *α* ∼ 0.4.

We finally settle on a network architecture that has one convolutional layer per block, skip connections, a dropout of 10% and a batch size of 50, while using the *Adadelta* optimizer. This network architecture has also been used for the training process shown in Fig. 3 and has been used to produce all results shown in the following. The entire network structure is implemented using the Keras Python API (41).

#### BFTTC algorithm

In order to evaluate the performance of our CNN in comparison to conventional TFM methods, we employ the Bayesian Fourier Transform Traction Cytometry (BFTTC) algorithm as a standard to compare with. The algorithm is described in Refs. (15, 30) and has been made publicly available by the authors at https://github.com/CellMicroMechanics/Easy-to-use_TFM_package.

## RESULTS

We analyze the performance of our ML approach on a set of error metrics. Additionally, we compare the performance to the BFTTC algorithm (15, 30) as a state-of-the-art conventional TFM method. Importantly, we want to discriminate between background noise and signal while also evaluating magnitude and angle reconstruction precision to infer whether our network generalizes to data never seen before. This will be done for synthetic displacement data first, which is generated in the same way as the training data and contains an additional varying level of noise. Subsequently, we can evaluate the performance of the CNN on artificial cell data using the same error metrics, on completely random displacement fields and, finally, we apply the CNN to real cell data.

### Evaluation metrics and application to synthetic patch-based data

We employ six evaluation metrics, four of which have already been introduced by Huang and coworkers (15) or earlier, to better distinguish between noise and signal, as well as between bias and resolution. Precise definitions of all evaluation metrics are given in the Appendix. In Fig. 5 we evaluate these metrics for varying noise levels during training of the network *N N*_low_ and in Fig. 6 we evaluate these metrics for varying noise levels 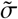 on our synthetic data patch-based data.

**Figure 5:**
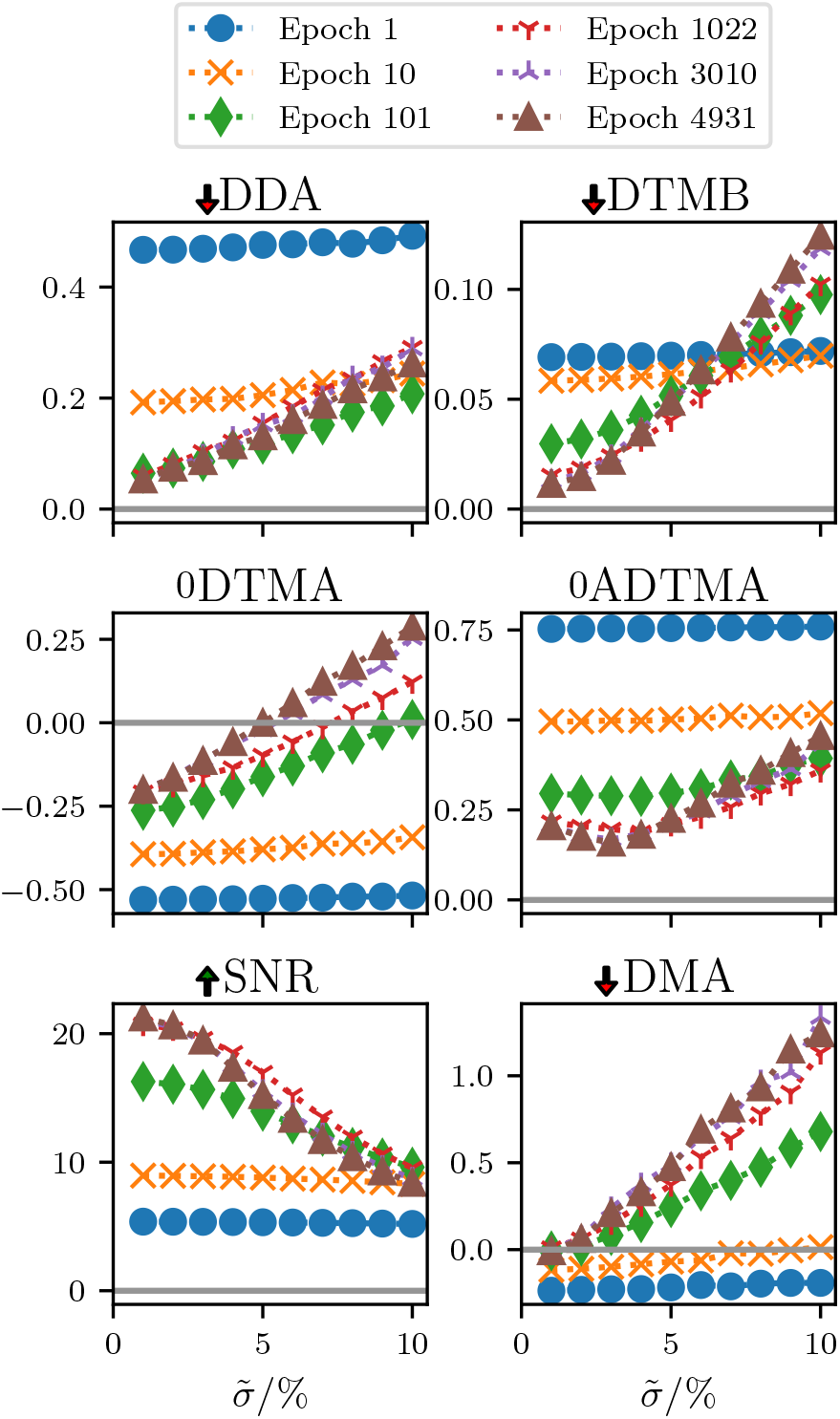
We evaluate the final low noise network *N N*_low_ during training by computing six precision metrics at intermediate points during training. We use the metrics to quantify the performance of the traction reconstruction during training of the network. This evaluation is performed with new data drawn from the same distribution and problem class as the training data. We clearly observe drastic improvements in predictive precision in the first 100 epochs, after which the improvement of the metrics is drastically slower. For the metrics DDA, DTMB and DMA a lower score is better (as indicated by the red arrow), while the SNR is better for larger values. Finally, the DTMA and ADTMA scores are better when they are closer to zero.

**Figure 6:**
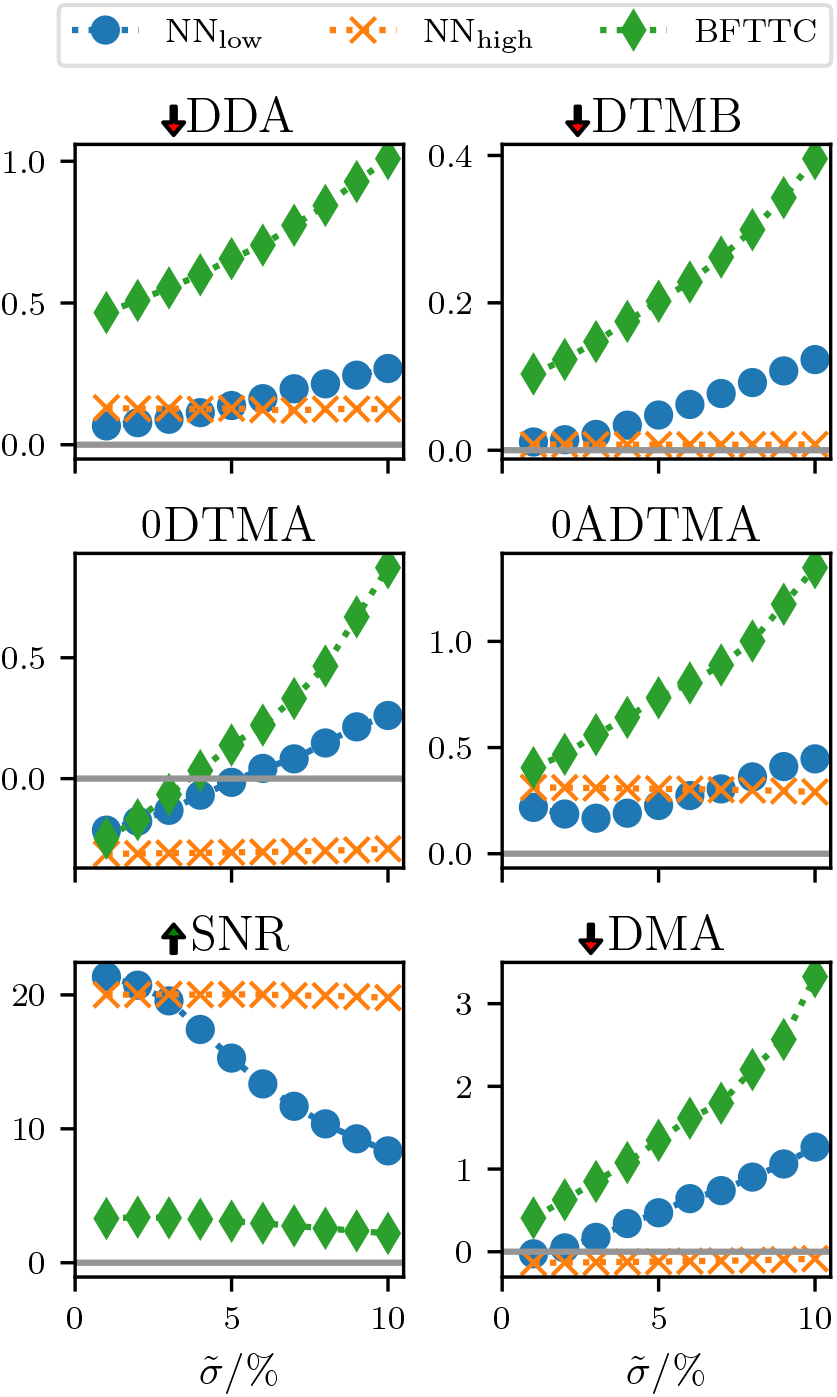
The comparison of our networks NN_low_ (trained with low noise background) and NN_high_ (trained with high noise background) with a state of the art conventional BFTTC approach shows the precision across the six evaluation metrics (see text) for varying noise levels 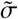 on synthetic patch-based data. This test is performed with an ensemble of traction spots randomly chosen in count, size, magnitude and orientation, testing our networks performance on data similar to the training data. The arrows next to the metric name indicate whether higher or lower is better; “0” indicates that the metric has a sign and a value of zero is optimal. Both of our networks outperform the BFTTC method in most metrics. With the high noise network NN_high_ we trade low noise fidelity for elevated noise handling capabilities.

The noise applied to the displacement field data is uncorrelated between pixels and randomly chosen from a Gaussian distribution centered around zero, with standard deviation σ. Let the dimensionless displacement field standard deviation be std*ū*, then we define our noise levels 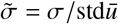, such that 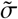 is the relative noise applied to the displacement field. In the following considerations we vary 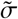 between 1 % and 10 %.

We pass the exact noise standard deviation to the BFTTC method for the noise evaluations, such that the BFTTC method has optimal conditions. Our networks do not get any additional information about standard deviation of the noise floor.

First, we introduce a measure to more precisely quantify the orientation resolution via the Deviation of Traction Direction at Adhesions (DDA)

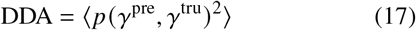

between predicted and true traction angles *γ* (see eqn. (28) in the Appendix for a more precise definition of the average); *p* (*α, β*) measures the *periodic* distance between two angles *α* and *β*. A small DDA indicates precise traction direction reconstruction. For both of our networks the direction reconstruction is more precise than the BFTTC method across the range of tested noise levels.

Second, we evaluate the Deviation of Traction Magnitude in the Background (DTMB) (15), which quantifies how accurate the traction magnitude reconstruction works in the background (see eqn. (24) in the Appendix), thus, if there is no prediction of an underground noise floor, not associated with any focal adhesion point, the DTMB score will be zero. Both of our NNs have a much lower DTMB score in than the BFTTC method in the limit 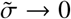, which should manifest in a much less noisy traction force reconstruction. While the low noise network again departs from that score linearly the high noise network again stays comparatively constant. During training we see that precision in low noise scenarios is traded for less robustness as evident from the increasing slope in Fig. 5. The high noise network again does not show this tendency.

Third, we discuss the Deviation of Traction Magnitude at Adhesions (DTMA) (13, 15) which evaluates the precision of traction magnitude reconstruction at the focal adhesion points (see eqn. (23) in the Appendix), thus the DTMA is zero for a perfect reconstruction, negative for an underestimation and positive for an overestimation of traction magnitudes. During training (see Fig. 5) this quantity consistently improves but trades precision in low noise scenarios for an increasing slope of the DTMA as a function of background noise. This is a first evidence that this network might perform poorly on high noise experimental data. We do not see the increase in slope for the high noise network (see Fig. 6). In Fig. 6 we see similar DTMA scores for all approaches in the limit 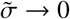, with a systematic under-prediction of tractions. For increasing noise floors both the BFTTC and the low noise network (NN_low_) depart from this common score and start to overestimate tractions. While the DTMA score for the high noise network barely changes, the low noise network DTMA score rises linearly with the noise floor 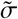, while the BFTTCs DTMA rises faster than linearly. The high noise network (NN_high_) retains a comparatively constant DTMA score and always under-predicts the traction magnitude.

Fourth, we introduce the Absolute Deviation of Traction Magnitude at Adhesions (ADTMA) that is similar to the DTMA, but evaluates the absolute deviations, capturing the actual reliability of reconstructions more precisely than DTMA, since alternating under- and over-predictions do not cancel out in this score. We can again see the same qualitative behavior as in most of the other metrics: Both networks are more precise in the low noise region, but the low noise network seems to perform best at a noise level of ∼ 3%, while the high noise network is robust against increases in background noise.

Fifth, the Signal to Noise Ratio (SNR) (15) also gives an insight into the noise floor of predictions (see eqn. (25) in the Appendix), it is high for a precise distinction between background noise and actual focal adhesion induced deformation and goes to zero for an increasingly noisy reconstruction. Both of our networks have a consistently higher SNR than the BFTTC method, undermining the assumption that the networks will yield a less noisy reconstruction overall. The low noise network SNR decays quickly with increasing noise levels, while the high noise network is more resilient against the increases in noise floor.

Finally, the Deviation of the Maximum traction at Adhesions (DMA) (15) gives a more detailed insight into the consistency for high amplitude tractions within a focal adhesion point (see eqn. (26) in the Appendix). A perfect reconstruction would yield a DMA score of zero, while under-predictions give negative scores and over-predictions positive scores. We can again observe that both of our networks give similar scores for low noise scenarios, which are both lower than the score of the BFTTC method. The low noise network again departs linearly from this common value, while the high noise network DMA score stays at a consistent level.

Across all six measures we observe the following trends for network NN_low_ trained with a low level of background noise as compared to network NN_high_ trained with a high level of background noise: NN_low_ perform superior to the BFTTC-standard *and* NN_high_ for low-noise data, because they are trained on low noise data. Their performance deteriorates, however, for higher noise levels 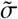, where their performance drops below NN_high_ but also below the BFTTC-standard. NN_high_ finds a better compromise between robustness and accuracy such that it outperforms the BFTTC-standard across *all* noise levels. Remarkably, the performance of the NN_high_ only deteriorates above noise levels 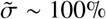.

### Tractions of artificial cells

In order to test the ability of the CNN to specialize from our general patch-based training set to realistic cell data, we first test the model on artificial cell data. The advantage of artificial cell data is that the *“true”* tractions are precisely known. A convenient model to generate realistic cell traction data artificially is the contractile network model (40). In this model, the stress fibers are active cable links with specific nodes anchored to the substrate at positions 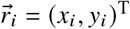. To construct a typical cell shape with lamellipodium and tail focal adhesions the tractions 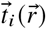 generated at each anchor point are generated by minimizing the total energy of the cable network. These tractions are then applied to circular patches of radius 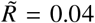 under an angle *γ*_*i*_ given by the stress fiber orientation at the anchored nodes (see Fig. 1).

In addition to Fig. 6, we want to evaluate the aforementioned metrics (DDA, ADTMA, DTMB, DMA, DTMA, SNR) on a displacement field generated by an artificial cell as shown in Fig. 1. Again, we add Gaussian noise to the displacement field with varying noise levels 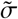 and evaluate the behavior of our networks NN_low_, NN_high_, and the BFTTC method in Fig. 7. We average all our results over 10 artificial cells.

**Figure 7:**
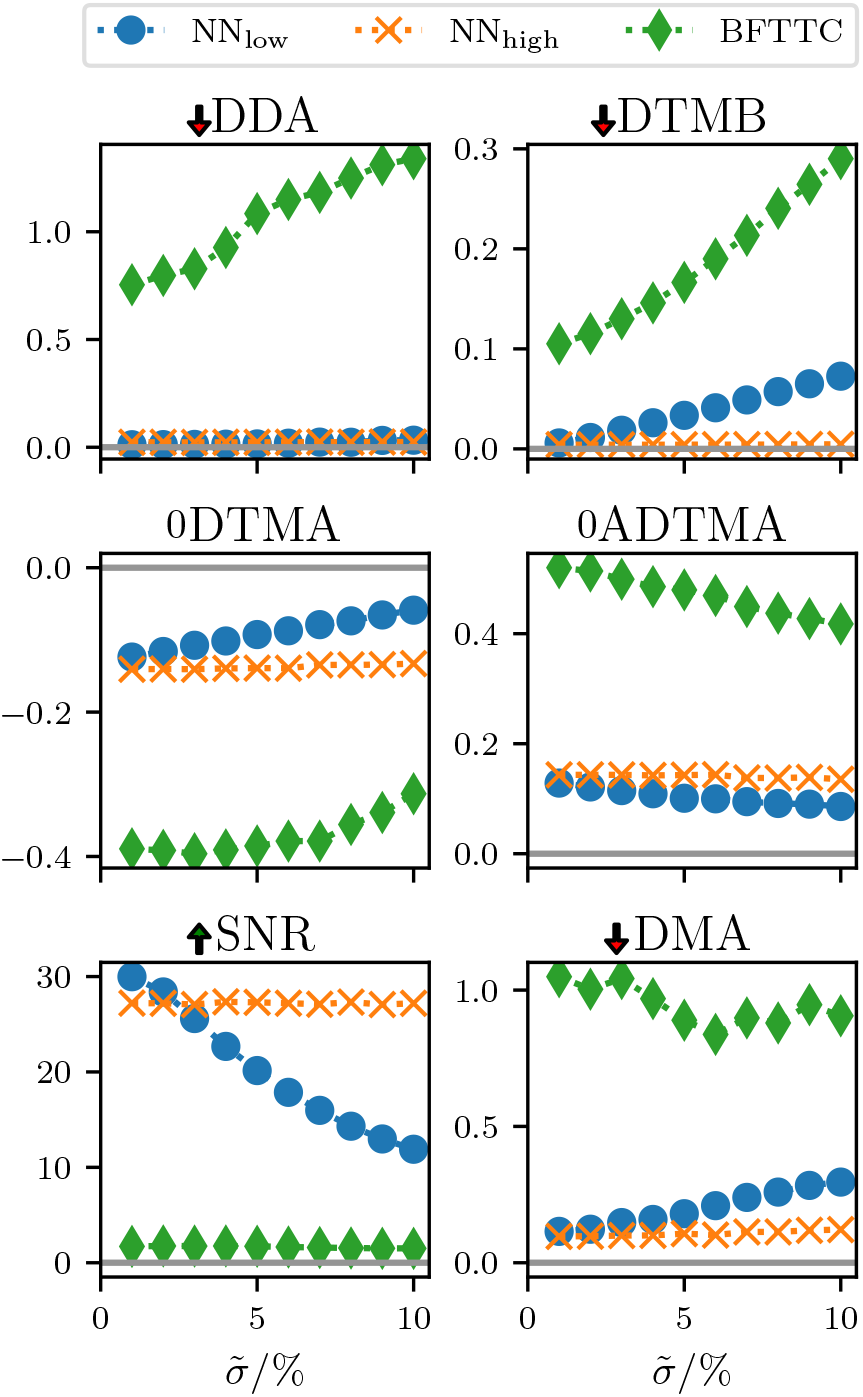
We compare our networks NN_low_ and NN_high_ with the BFTTC method for data generated from an artificial cell model using the same six evaluation metrics for varying noise levels 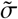 as in Fig. 6. Since the method of generating the data is now different from the training process, we expect to see the generalization potential of our networks more clearly than in Fig. 6. The BFTTC method works considerably worse on this data compared to Fig. 6. Additionally, we see an amplified underestimation of the traction magnitudes for the BFTTC method. With respect to noise, the high noise network seems to give the best compromise between precision and regularization of the output traction fields.

We see qualitatively similar results to Fig. 6 in the SNR and DTMB metrics, while the absolute performance in those metrics is better (higher SNR and lower DTMB scores) for the artificial cell data. This is likely due to the lower number of traction patches in total and the equal radii of all traction patches. Both networks, are able to reconstruct the traction fields more reliably and with greater precision. This is true across all observed metrics.

The artificial cell data show a clear tendency towards a traction magnitude underestimation (DTMA, ADTMA) for all approaches. Since we are generating tractions in strongly bounded range due to the cable network, the traction spots tend to be in close proximity to each other, which can increase smearing of sharply separated traction spots.

### Reconstruction of random traction fields

In order to prove that our networks indeed have learned to exploit the linearity of the problem and that they have learned a general solution for the problem, we probe them on entirely random traction fields. These are generated starting from spatially uncorrelated Gaussian noise, which is subsequently convolved with a proximity filter with a characteristic correlation length of ∼ 0.1 in dimensionless units of the image size. Then we compute the corresponding displacement field and pass it to our high noise network and the BFTTC method for reconstruction.

The advantage of this reconstruction setting is that we have the exact ground truth for tractions while not using traction fields that are similar to the training data. As detailed in the Supporting Material, our high noise network can reconstruct the random traction field and associated displacements with higher accuracy than the BFTTC method.

### Tractions of real cells

Finally, we want to test our ML approach on real cell images. Of course, we do not have access to the *“true”* traction field for those images, however, we can qualitatively compare the results obtained from our ML approach with those obtained by the well tested BFTTC method. The example cell results shown in Fig. 8 are for a NIH/3T3 (National Institutes of Health 3T3 cultivated) Fibroblast on a substrate with length *L* = 200.2 *μ*m and elastic modulus *E* = 10670 Pa (the data was made available in Ref. (17)). Additional results for all 14 cells provided in Ref. (17) are similar and provided in the Supporting Material.

**Figure 8:**
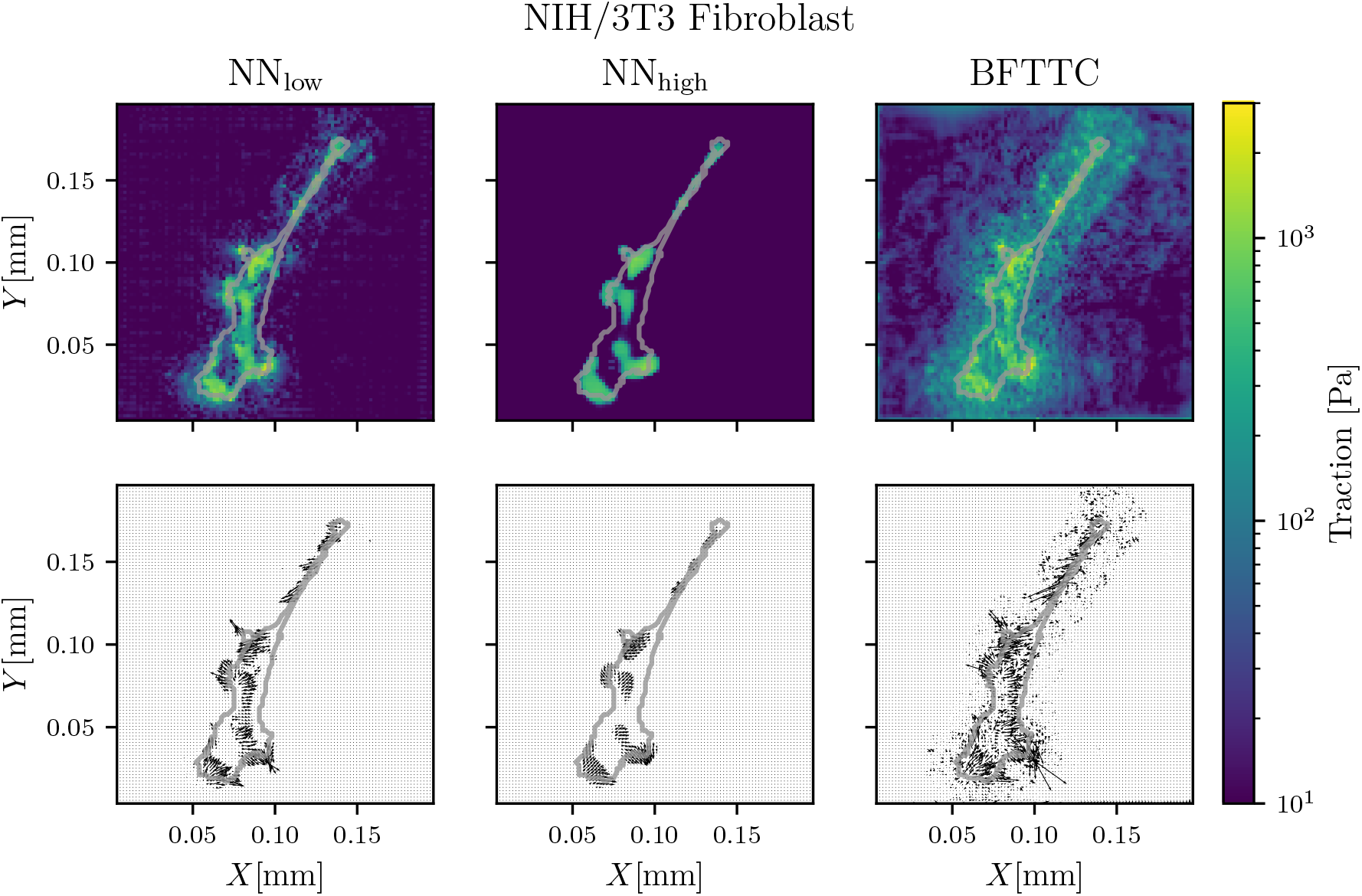
Comparison of traction reconstruction for a real Fibroblast between our networks NN_low_ and NN_high_ and the BFTTC method (top and middle row). Although we do not have the “true” traction field at hand for a quantitative evaluation of precision across the methods, we see compatible results across the board, while both networks have a significantly reduced noise floor. The top row shows traction magnitude reconstruction, while the center row shows angle reconstruction.

It is apparent that the network trained with low noise reconstructs a traction field, which is similar to that of the BFTTC method, while the noise in the vicinity of the cell is significantly reduced. The network trained with a high noise floor gives a more regular traction pattern and cuts off lower amplitude tractions.

The reason for these results is that the network trained with low noise exhibits a SNR superior both to BFTTC and the network trained with high noise levels if noise in the experimental data is low (see Figs. 6 and 7); for artificial cell data also the DTMA and ADTMA of the network trained with low noise is superior for low experimental noise levels (see Fig. 7). A low noise in experimental data seems to be realized here. We can thus infer that the tractions reconstructed by the high noise network systematically under-predict the real tractions for this particular data.

As expected from our prior analysis, the resistance to additional noise is much better for the network which saw high noise levels during training and the low noise network is fails completely when subjected to very high noise. The robustness of the high noise network is highlighted in Fig. 9, where the cell data is superposed with significant background noise of 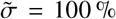. Both our low noise network and the BFTTC method produce a noisy traction field in this case, while the high noise network still displays a similar traction pattern in both of these cases. The noise seen in the BFTTC method is significantly reduced, as we pass the *exact* standard deviation of the applied noise to the method.

**Figure 9:**
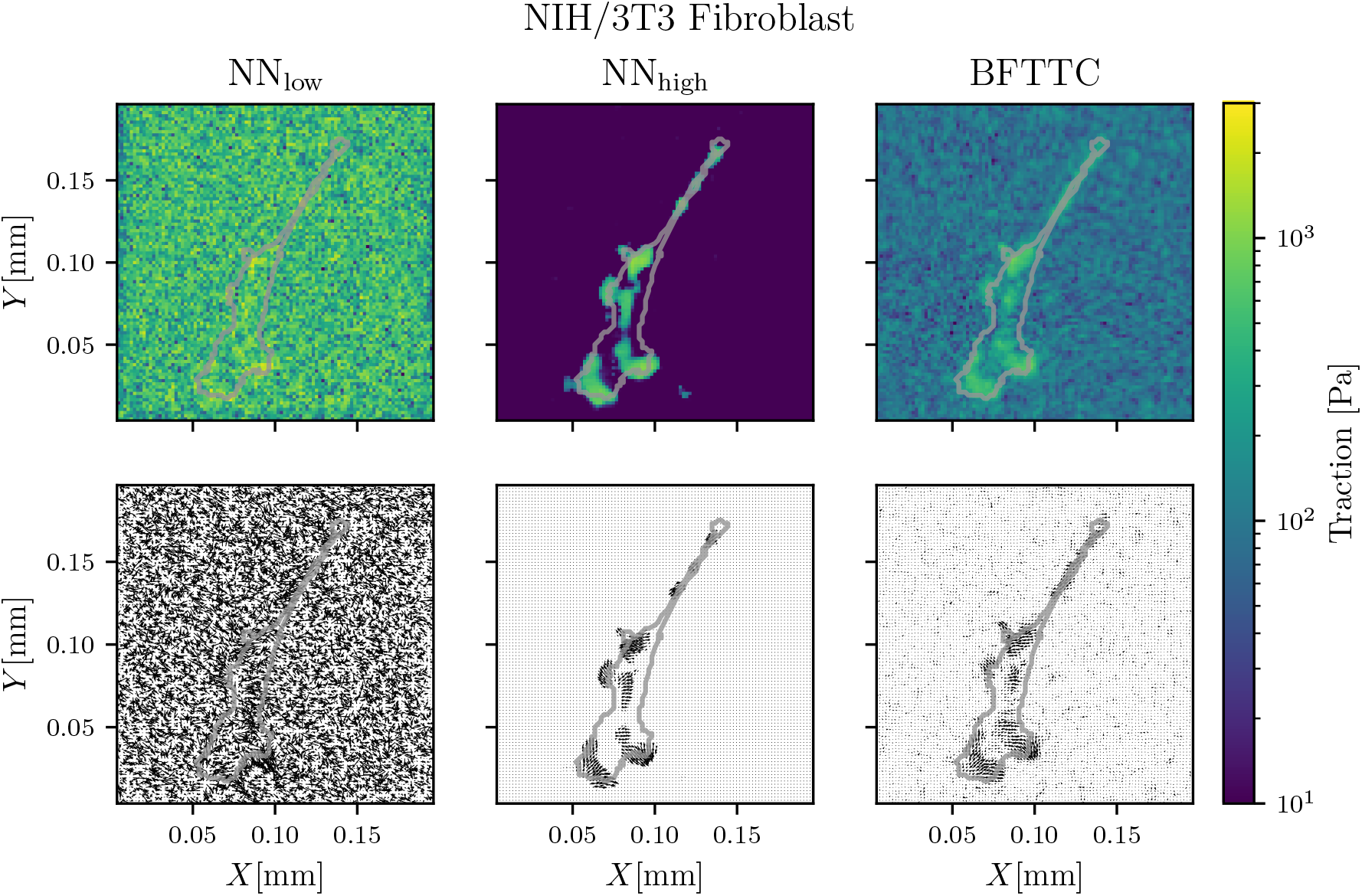
Adding noise to the Fibroblast displacement field shows strong noise robustness of our network NN_high_, which has been trained for high noise scenarios, in traction reconstruction (top and middle row). The low noise network NN_low_ fails to compensate for the high noise and the BFTTC method yields qualitatively similar results to the high noise network, but exhibits strong noise artifacts in the reconstructed traction fields. The BFTTC method has an unfair advantage, since we pass the *exact* standard deviation of the perfectly uniform Gaussian noise to it for its regularization, while our networks do not get this information. The bottom row displays the displacement field computed from the reconstructed traction fields.

Overall, the result from the high noise network and the BFTTC method is qualitatively similar in the high noise scenario, while the high noise network reconstruction is more regular and fits the circular focal adhesion point model more consistently and stays invariant for a wide range of background noise levels. Apparently, the noise the network sees in training directly controls the regularization of the reconstruction and it would be possible to create intermediate networks with higher sensitivity, but lower noise invariance.

Finally, we compute the residual error between the experimental (already noisy) input image and the reconstructed displacement field, by solving the reconstructed traction field for the displacement field. We do this for all the 14 cells provided in Ref. (17) (see Supporting Material). For the low noise network we achieve a mean RMSE ∼ 10^−3^, while the high noise network performs slightly better at an RMSE ∼ 9 · 10^−4^. The BFTTC method performs significantly better with an average RMSE ∼ 5 · 10^−5^. While the residual errors are low for all approaches we can conclude that the BFTTC method reconstructs the input displacement field more accurately. This seems surprising in light of the higher traction background noise outside the cell shape that the BFTTC clearly produces. The reasons are discussed in the next section in more detail.

We can additionally quantify the contribution of tractions that lie outside the cell contour, which can be considered unphysical. For this we subtract all tractions inside of the contour from the full traction fields and are left with tractions that lie outside the cell contour. When calculating the rooted mean square of these outside tractions we are left with a metric that quantifies physical consistency. We perform this analysis for the cell in Fig. 8 and find that the mean background traction for the BFTTC method is ∼ 0.013, while it is ∼ 0.009 for NN_low_ and ∼ 0.008 for NN_high_. Since the cell segmentation is only an approximation this metric is only a proxy for physical consistency.

We conclude that the high noise network is the better choice for a traction field where the noise is not known, or might be inhomogeneous. If the experimental error is high, or the displacement field reconstruction is imprecise the high noise network provides a robust way of extracting the traction field, while conventional methods are plagued by high background noise in the reconstructed traction fields in this case.

Evaluating a traction field with our networks takes 1 ms, while evaluation an image with the BFTTC method takes 1.5 s (with size 104 × 104). This is a performance improvement of more than three orders of magnitude. Because of the nondimensionalization we perform it is only necessary to train one network for a large range of experimental realizations. Thus, when the same network is reused for a number of different experiments the long training time will eventually be outweighed by the significant performance advantage at inference.

### Displacement versus traction error

We train our NNs for correct tractions by using the MSE of tractions from eq. (16) as objective function. This is only possible with synthetic data, where the “truth” for tractions is known. Within this approach we optimize accuracy in tractions but concede errors in the displacement which are not of primary interest in TFM.

Alternatively, one could base training on the MSE of displacements with the drawback of slowing down the training process by orders of magnitude but with the advantage that training could also be performed with actual experimental data. Networks trained with displacement-based metrics will minimize the displacement error in order to obtain correct tractions, which is a more indirect approach. The BFTTC algorithm also minimizes the deviations in displacements in order to determine an optimal traction field (15). We show that this causes deviations in tractions by applying both strategies to situations where we know the “true” tractions.

Figure 10 shows a comparison for a synthetic traction field that we generate from the NIH/3T3 traction data (see Fig. 8) by suppressing tractions outside the cell shape (which should be artifacts). This provides the traction “truth” for the comparison. The corresponding displacement field is computed and spatially uncorrelated low amplitude Gaussian noise is added. This displacement field is analyzed by the low and high noise networks and the BFTTC algorithm. Fig. 10 clearly shows that, on the one hand, both low and high noise network give a significantly better traction reconstruction, in particular outside the cell shape where the BFTTC method tends to generate background traction noise. On the other hand, this background traction noise obviously enables the BFTTC method to lower the displacement errors, in particular inside the cell shape. As we are interested in correct traction reconstruction in TFM, this comparison clearly pinpoints the advantages of the CNN reconstruction when trained with the traction MSE as objective function.

**Figure 10:**
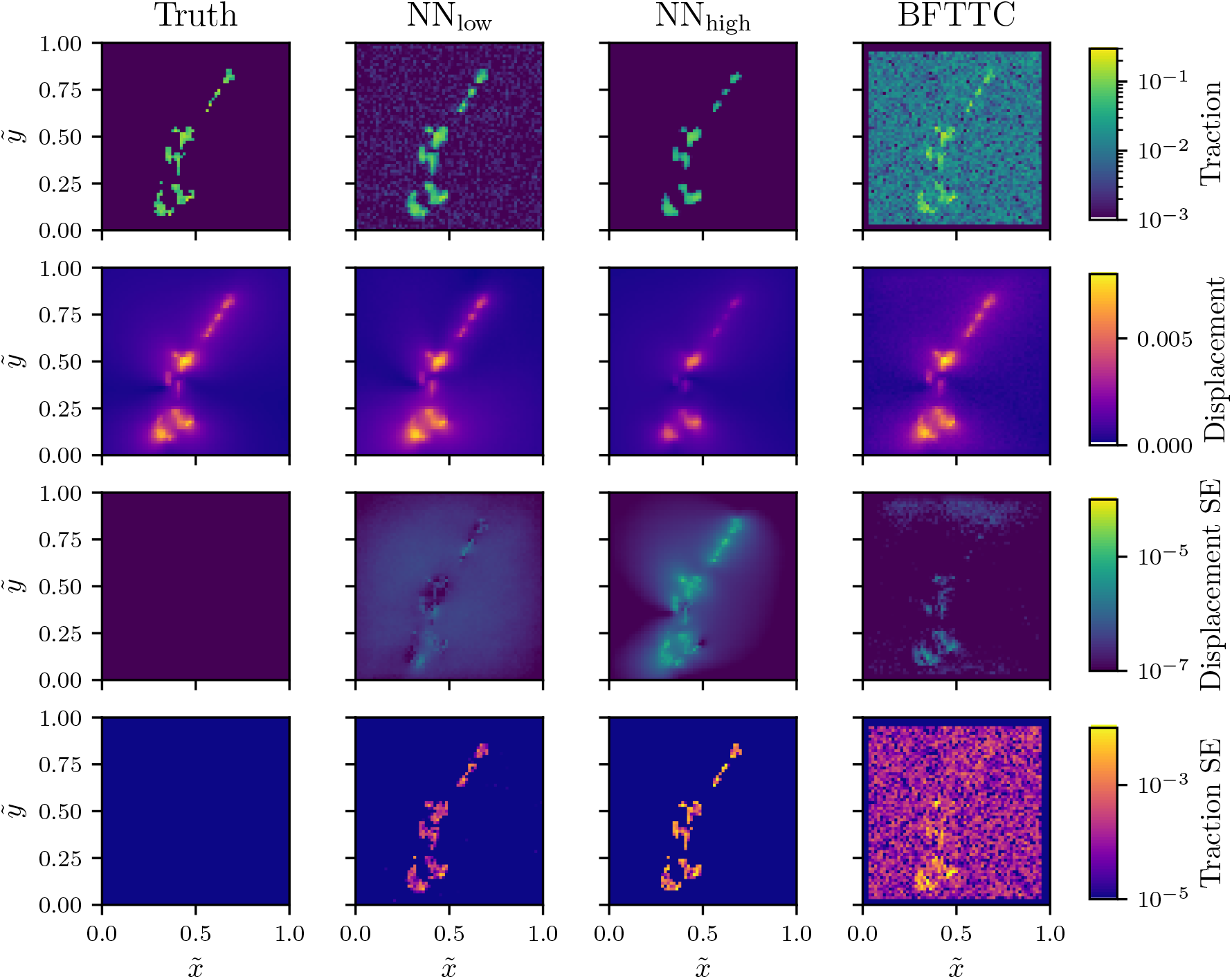
Comparison of traction and displacement error between our networks NN_low_ and NN_high_ and the BFTTC method for a synthetic Fibroblast-like traction field and corresponding calculated displacements. The two top rows show the reconstructed traction and displacement fields as compared to the “truth” (first column), which is at hand in this comparison. The two bottom rows display the square error calculated between the “true” and reconstructed displacement and traction fields. We see that both NNs give better traction reconstruction with reduced noise outside the cell as compared to BFTTC, while displacement errors are slightly higher.

### Input resolution scaling

So far we used a fixed size of *N* × *N* = 104 × 104 for input images. Since our networks are exclusively composed out of input size independent layers, we are able to feed arbitrary input sizes (i.e., arbitrary image resolutions) to the networks.

It is, however, important to realize that we provide no spatial information to the network apart from the size *N* × *N* of the input array. Because the total input size is not visible to our convolutional layers the network has no means to adapt to changes of the input size *N*. It is possible to circumvent this limitation by scaling the input displacements properly, such that the input is locally equivalent to that of a 104 × 104 grid. A local 104 × 104 section of a *N* × *N* image of a substrate of length *L* corresponds to a section of smaller length (104/*N*) *L* such that the dimensionless displacement *ū* for a resolution *N* × *N* corresponds to a larger dimensionless displacement (*N*/104) *ū* for a 104 × 104 section of the same substrate. Or, in other words, the displacement scale must be coupled to the pixel scale, since our networks directly operate on the pixel level with dimensionless displacements. The scale factor for the input displacements for an image of size *N* × *N* is thus *N*/104. We show in the Supporting Material that this is sufficient to make the networks usable for arbitrary resolutions. We compare the performance of our networks with the BFTTC method on 256 × 256 grids and find superior precision and speed provided by our networks. In particular, the SNR metric further improves upon increasing resolution.

## DISCUSSION

We present a ML approach to TFM via a deep convolutional NN trained on a general set of synthetic displacement-traction data derived from the analytic solution of the elastic forward problem for random ensembles of circular traction patches. This follows the general strategy that NNs trained with easy-to-generate data of representative forward solutions can serve as a regressor to solve the inverse problem with high accuracy and robustness.

Our approach to TFM uses synthetic training data derived from superpositions of known and representative traction patches. This allows us to employ an objective training function that directly measures traction errors. This contrasts conventional TFM approaches such as BFTTC, where the tractions are adjusted to match the displacement field (eventually subject to additional regularizing constraints on the tractions), such that low displacement errors are the implicit objective. We show that a force conserving discretization is crucial for high performance networks and we find a significant enhancement of the robustness of the NN if the training data is subjected to an appropriate level of additional noise.

In conventional TFM approaches the inverse elastic problem is ill-posed and the suitable choice of regularization in the inversion procedure is crucial and has been a topic of active research over the last twenty years. ML approaches circumvent the need for explicit regularization and provide an implicit regularization by a proper choice of the network architecture, i.e., convolutional NNs for TFM, and after proper training. Our work shows that the suitable choice of “physics-informed” training data and, moreover, the suitable choice of noise on the training data governs the applicability of the NN and the compromise between accuracy and robustness in ML approaches, somewhat analogous to the role of the regularization procedure in conventional TFM approaches.

We employ a sufficiently general patch-based training set and show that this allows the CNN to successfully specialize to artificial cell data and real cell data. Moreover, training with an additional background noise that is 5% of the average variance of the dimensionless displacement field (the *N N*_high_ network eqn. (15)), gives a robustness against noise in the NN performance that is superior to state-of-the-art conventional TFM approaches without significantly compromising accuracy. We can systematically back these claims by characterizing both the NN performance and the performance of state-of-the-art conventional TFM (the BFTTC method) via six error metrics both for the patch-based training set (Fig. 6) and the artificial cell data set (Fig. 7), which are two data sets where we can compare the prediction to the true traction labels. We also test the NN performance on random traction fields (see Supporting Material) and traction fields derived from real cell data (Fig. 10). Whenever the true tractions are known, we find that our NNs, which were trained to minimize traction errors, give more accurate traction reconstruction with a reduced background traction noise outside the cell shape, although the NNs tend to concede higher errors in the corresponding displacement fields.

For real cell data, we find that a NN trained with low noise (0.5%) gives the best performance if the experimental data is of high quality with low noise levels (see Fig. 8). For noisy experimental data, on the other hand, the NN trained with high levels of noise (5%) clearly performs best (see Fig. 9). This suggests that it might be beneficial to first employ the high noise network on experimental data and only switch to the low noise network if the background noise level is below 1% of the displacement standard deviation.

Overall, we provide a computationally efficient way to accelerate TFM as a method and improve both on accuracy and noise resilience of conventional approaches, while reducing the computational complexity, and thus execution time by multiple orders of magnitude compared to state-of-the-art conventional approaches. It is apparent from our analysis that ML approaches have the potential to shift the paradigm in solving inverse problems away from conventional iterative methods, towards educated regressors which are trained on a well understood and numerically simple to solve forward problem.

We make all NNs discussed in this work freely available for further use in TFM. We use a 104 × 104-grid for the displacement data, but show that our networks are able to handle arbitrary displacement data resolutions. Experimental data can easily be adapted to comply with the network input shape by properly scaling the displacements or, alternatively, by interpolating or downsampling to a 104 × 104-grid, which will, however, decrease the traction resolution.

By using non-dimensionalized units, the NNs made available with this work are widely applicable across different problems and can also be easily further adapted, for example, to problems where typical tractions are not limited to the range traction ranges discussed here by repeating the training process. In the Supporting Material, we show that the present networks are able to generalize to larger dimensionless traction magnitudes than trained for, with 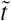 ranging up to 1.5, without re-training. Another potential problem to be addressed in future work is the effect of spatial noise correlations, for example, from optical aberration or from the displacement tracking routine that is applied to generate the displacement input data. In the Supporting Material, we consider uncorrelated Gaussian noise with a standard deviation that decreases with the distance from the image center and find a robust performance of the high noise network. Robustness to noise with genuine spatial correlations over characteristic distances significantly larger than the pixel size will presumably require re-training of the networks. All necessary routines to re-train a NN to new traction levels, new characteristic patch sizes, or other noise levels are made freely available with this work at https://gitlab.tu-dortmund.de/cmt/kierfeld/mltfm. This will also allow to easily adapt the training to other types of noise correlations.

## AUTHOR CONTRIBUTIONS

F.S.K, L.M. and J.K. developed the theoretical and computational model. F.S.K. and L.M. performed all computations, construction and training of neural networks, and data analysis. F.S.K and J.K. wrote the original draft. F.S.K., L.M. and J.K. reviewed and edited the manuscript.

## ACKNOWLEDGMENTS

F.S.K. acknowledges support by the German Academic Scholarship Foundation.

## SUPPORTING MATERIAL

Supporting Material can be found online at http://www.biophysj.org.

## SUPPORTING CITATIONS

References 11, 17, 27 appear in the Supporting material.

## DETAILS OF THE SOLUTION OF THE FORWARD PROBLEM FOR TRACTION PATCHES

For the solution of the forward problem for a tractions patch of size *R*_*i*_, we obtained the functions 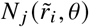 in eqn. (12) and eqn. (13). They are defined as

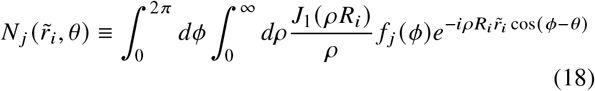

with

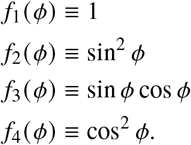

By definition, *N*_2_ + *N*_4_ = *N*_1_.

The remaining integrals in the functions 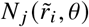 can be performed analytically:

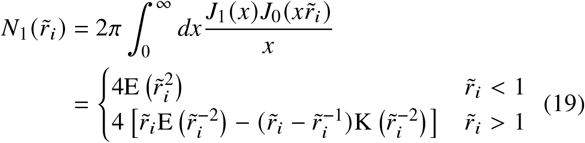

with the complete elliptic integrals 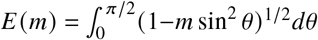 and 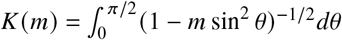;

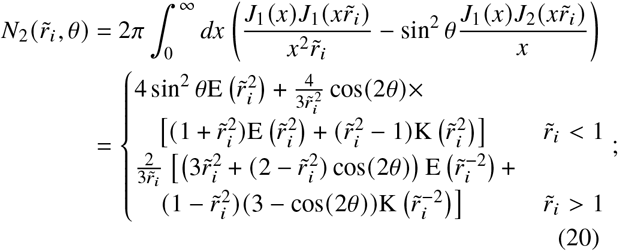

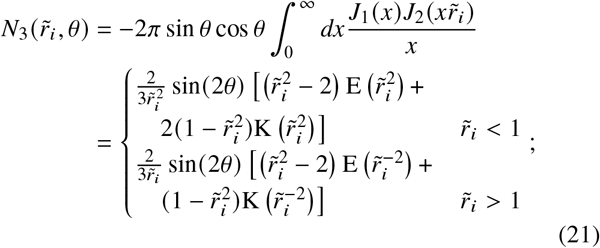

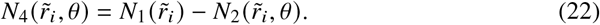

## DEFINITIONS OF EVALUATION METRICS

We employ six evaluation metrics (see Figs. 6 and 7). Their definition is based on a comparison of traction predictions 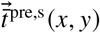 in sample s compared to “true” tractions 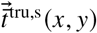, which are known for the artificial data for random circular traction patches. We evaluate all six metrics by averaging over *S* = 100 samples; the sample average is denoted by ⟨… ⟩. All traction vectors 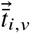 in patch *i* (*i* = 1, …, *n*) are indexed by *v*. All tractions vectors 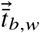 outside patches are considered as belonging to the background *b* and indexed by *w*. For completeness we give the precise definitions of all six evaluation metrics:

1. Deviation of Traction Magnitude at Adhesions (DTMA) (15):

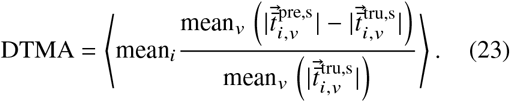

Note that mean 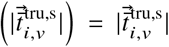 because artificial traction data is piecewise constant in traction patches.
2. Deviation of Traction Magnitude in the Background (DTMB) (15):

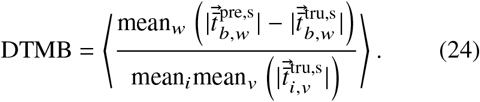

Note that 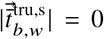 because artificial traction data exactly vanishes outside patches.
3. Signal to Noise Ratio (SNR) (15):

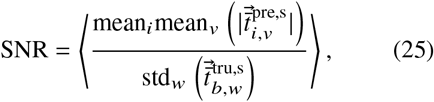

where std is the standard deviation.
4. Deviation of the Maximum traction at Adhesions (DMA) (15):

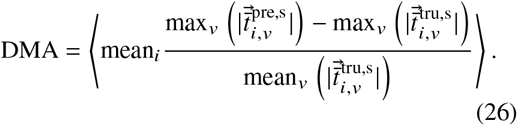

Note that max 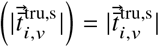 because artificial traction data is piecewise constant in traction patches.
5. Absolute Deviation of Traction Magnitude at Adhesions (ADTMA):

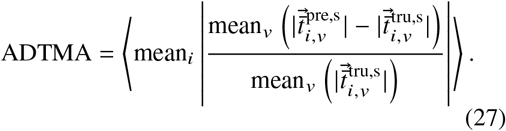
6. Deviation of Traction Direction at Adhesions (DDA):

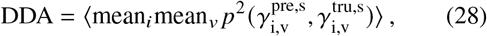

where *p* (*α, β*) measures the *periodic* distance between two angles *α* and *β*.

