## Supporting Material for "Enhancing Robustness, Precision and Speed of Traction Force Microscopy with Machine Learning"

### REDUCTION OF THE ELASTIC GREEN'S TENSOR TO TWO DIMENSIONS

The full three-dimensional Green's tensor of the boundary problem in the main text on the half-space $z > 0$ is given by [S1, S2]:

$$\mathbf{G}(x,y,z) = \frac{1+\nu}{2\pi E}\begin{pmatrix} \frac{2(1-\nu)r+z}{r(r+z)} + \frac{(2r(\nu r+z)+z^2)x^2}{r^3(r+z)^2} & \frac{(2r(\nu r+z)+z^2)xy}{r^3(r+z)^2} & \frac{xz}{r^3} - \frac{(1-2\nu)x}{r(r+z)} \\ \frac{(2r(\nu r+z)+z^2)xy^2}{r^3(r+z)^2} & \frac{2(1-\nu)r+z}{r(r+z)} + \frac{(2r(\nu r+z)+z^2)y^2}{r^3(r+z)^2} & \frac{yz}{r^3} - \frac{(1-2\nu)y}{r(r+z)} \\ \frac{xz}{r^3} + \frac{(1-2\nu)x}{r(r+z)} & \frac{yz}{r^3} + \frac{(1-2\nu)y}{r(r+z)} & \frac{2(1-\nu)}{r} + \frac{z^2}{r^3} \end{pmatrix}, \tag{S1}$$

The tensor is not symmetric, in general. On the surface $z = 0$, this becomes

$$\mathbf{G}(x,y,0) = \frac{1+\nu}{\pi E r^3}\begin{pmatrix} (1-\nu)r^2 + \nu x^2 & \nu xy & -\frac{1}{2}(1-2\nu)xr \\ \nu xy & (1-\nu)r^2 + \nu y^2 & -\frac{1}{2}(1-2\nu)yr \\ \frac{1}{2}(1-2\nu)xr & \frac{1}{2}(1-2\nu)yr & (1-\nu)r^2 \end{pmatrix}. \tag{S2}$$

For incompressible polymer materials with $\nu \approx 1/2$, the coupling between in-plane tractions and out-of-plane displacements is small.

### DISCRETIZATION OF TRACTION PATCHES

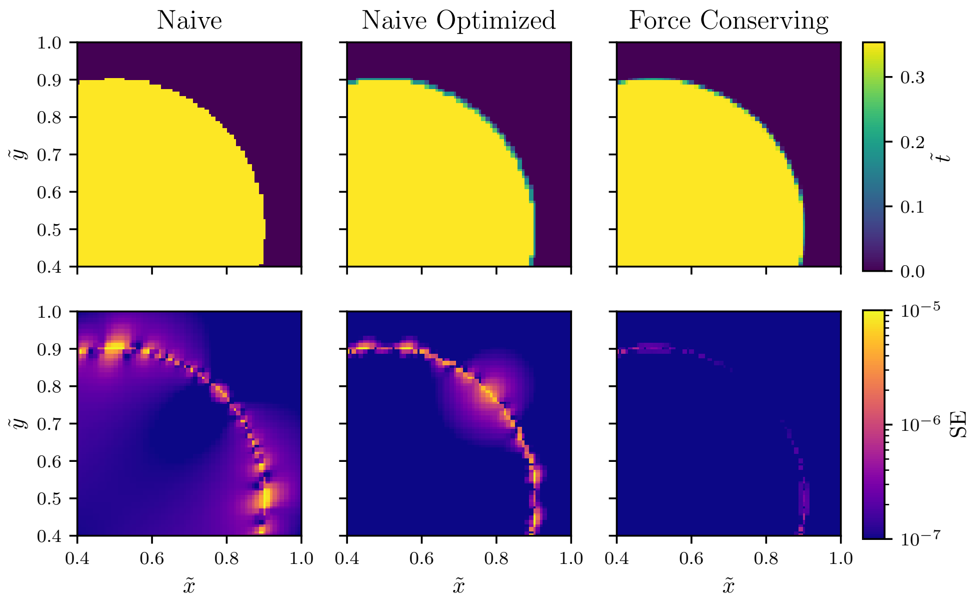


FIG. S1. The discretization of circular traction patches onto a pixel grid (top row) with a naive discretization, an improved discretization and a force conserving discretization (left to right). The resulting residual displacement errors (bottom row) are generated by feeding the discretized traction field back to the solver to generate a residual of the input displacement field and the displacement field obtained from resolving the discretized traction field. The residual error decreases drastically if the total traction force is conserved by the discretization method.

A naive approach for the discretization of the circular traction patches onto a square pixel grid with indices $i, j \in \{1, ..., N\}$ and square side lengths $a = L/N$ is to check whether any square segment center point $\vec{c}_{ij}$ is contained in the circular traction patch of radius $R_t$ and center point $\vec{c}_t$. If the center point is contained, the grid segment $ij$ is assigned the traction $\tilde{t}_0$ of the circular patch:

$$\tilde{t}_{ij} = \begin{cases} \tilde{t}_0, & \text{for } |\vec{c}_{ij} - \vec{c}_t| < R_t \\ 0, & \text{else.} \end{cases} \tag{S3}$$

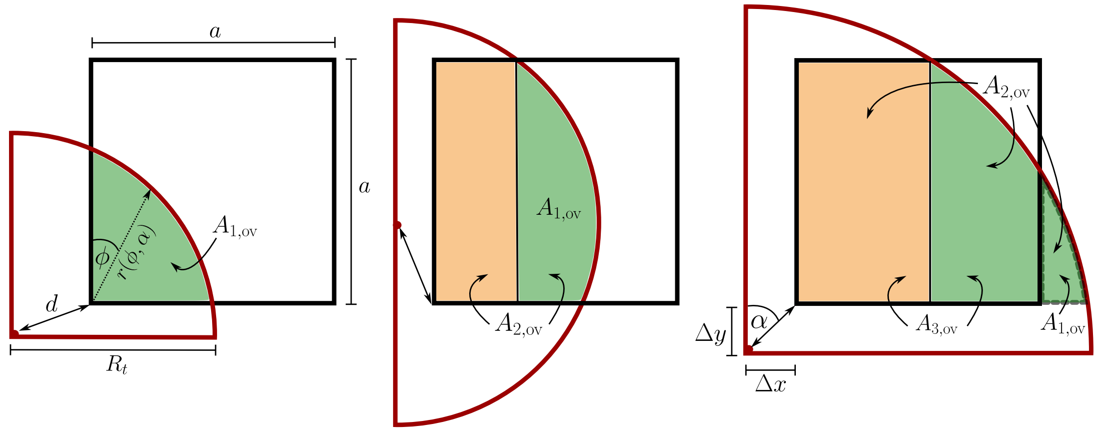


FIG. S2. Calculation of overlap areas $A_{k,\mathrm{ov}}$ for circular patches containing $k = 1, 2, 3$ corners of square grid segments.

This naive discretization suffers from a critical artifact: it does not conserve the total traction force $F_t$ exerted by the patch, $\sum_{i,j} a^2 \tilde{t}_{ij} \neq \tilde{t}_0 \pi R_t^2 = F_t$.

One step towards improving the force conservation is to check for all four corners of the square grid segment whether they are contained in the traction circle. If we assign equal weights to any of the corners,

$$\tilde{t}_{ij} = t_0 \frac{\#\text{corners contained}}{4}, \tag{S4}$$

the naive optimization is improved, but still not force conserving.

Ultimately, we construct a force conserving discretization by explicitly calculating the *exact* area occupation ratio of each square grid segment and the circular traction spots (see Fig. S2).

We start with the case that exactly one corner of the grid square with is contained in the traction circle. We only need to consider the case where only the bottom left corner of the grid segment is contained in the traction circle; all remaining cases can be constructed by mirror or rotational symmetries. We denote the vector from the center of the circle to the bottom left corner of the grid segment by $(\Delta x, \Delta y)$, its length by $d$ and the angle with the y-axis by $\alpha$ such that $(\Delta x, \Delta y) = d(\sin\alpha, \cos\alpha)$. The overlap area of traction circle and grid square is given by the polar integral

$$A_{1,\mathrm{ov}} = \frac{1}{2}\int_0^{\pi/2} \mathrm{d}\phi r^2(\phi,\alpha) \tag{S5}$$

with

$$r(\phi,\alpha) = \sqrt{R_t^2 - d^2\sin(\phi-\alpha)} - d\cos(\phi-\alpha)\,. \tag{S6}$$

This integral can be solved via basic calculus:

$$\begin{aligned} A_{1,\mathrm{ov}}(\Delta x, \Delta y) = {} & \frac{1}{4}R_t^2 + \Delta x \Delta y \\ & - \frac{1}{2}\left(\sqrt{R_t^2 - \Delta y^2}\Delta y + \sqrt{R_t^2 - \Delta x^2}\Delta x + R^2 \arcsin(\Delta y/R_t) + R^2\arcsin(\Delta x/R_t)\right)\,. \end{aligned} \tag{S7}$$

If two corners of the grid square are contained in the traction patch it suffices to consider the case where the bottom left and the top left corner are contained because of rotational symmetries. Additionally, we can concentrate on the case where the center point of the traction patch is *below* (smaller y-component) the grid square center point because of the mirror symmetries of the problem. The two corner overlap can be viewed as a one corner overlap with a shifted $\Delta x$ and an additional rectangular overlap (see Fig. S2) resulting in

$$A_{2,\mathrm{ov}}(\Delta x, \Delta y) = A_{1,\mathrm{ov}}\left(\sqrt{R_t^2 - (\Delta y + a)^2}, \Delta y\right) + a\left(\sqrt{R_t^2 - (\Delta y + a)^2} - \Delta x\right)\,. \tag{S8}$$

Similarly, the three corner overlap decays into a two corner overlap minus a shifted one corner overlap (see Fig. S2). Again, for symmetry reasons, we only need to consider one configuration, where the top left, bottom left and bottom right corner of the grid square are contained in the circular traction patch:

$$A_{3,\mathrm{ov}}(\Delta x, \Delta y) = A_{2,\mathrm{ov}}\left(\Delta x, \Delta y\right) - A_{1,\mathrm{ov}}\left(\Delta x + a, \Delta y\right)\,. \tag{S9}$$

Finally, in order to conserve the total force exerted by the traction patch, the tractions assigned to a grid square must then be weighted by their area occupation percentage,

$$\tilde{t}_{ij} = \tilde{t}_0 \frac{A_{\mathrm{ov},ij}}{a^2}\,, \tag{S10}$$

where $A_{\mathrm{ov},ij}$ is calculated from eqns. (S7), (S8), or (S9) depending on the number of corners of square $ij$ contained in the circular traction patch.

We integrate this force balance conserving discretization algorithm in a solver to generate the displacement fields from arbitrary superpositions of circular traction patches. The solver is implemented in a C/C++ package with a Python binary interface to combine the speed of native byte code with the simplicity of an interpreted language.

The fundamental error of this method can be computed by generating a large exact circular solution and feeding the discretized traction grid back into our solver. With the force conserving implementation we are able to reproduce the displacement field of a large circular traction patch of radius $R_t = 0.4$ to a precision of RMSE $\sim 1.2 \cdot 10^{-4}$. This also allows us to probe the difference between the three traction discretization methods in Fig. S1. The naive discretization gives a residual error RMSE $\sim 5.1 \cdot 10^{-4}$ for the displacement field, the optimized naive discretization is already able to lower the residual error drastically to RMSE $\sim 4.4 \cdot 10^{-4}$. The force conserving method finally lowers the residual error to RMSE $\sim 1.2 \cdot 10^{-4}$, which is significantly better than both the naive and the improved naive method.

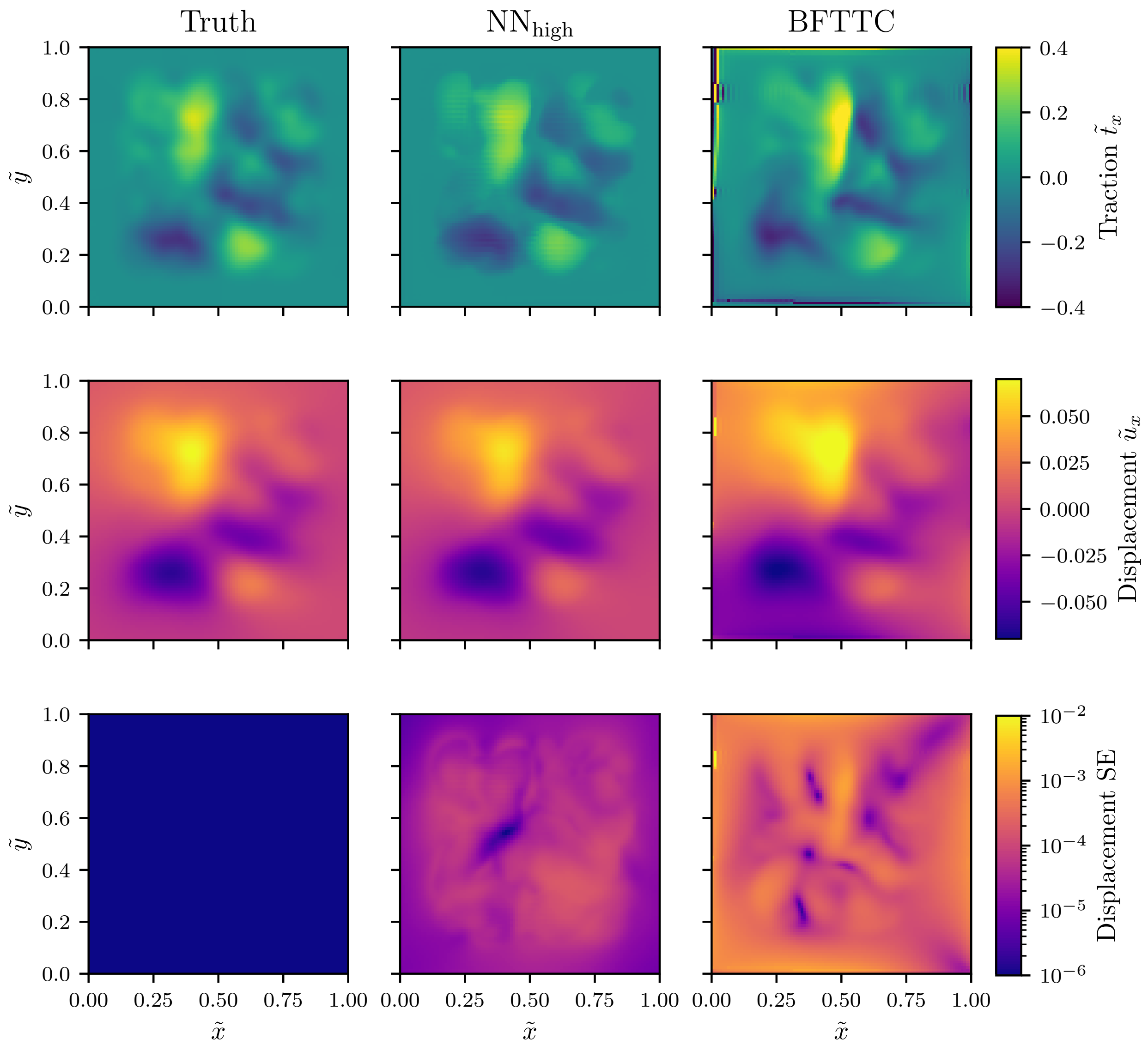


FIG. S3. We construct a completely random traction field that is generated by an entirely different method, as compared to the training data. From the generated random traction field (upper left) we generate the accompanying displacement field (middle left), which we feed to our high noise network and the BFTTC method. Both methods generate a traction field reconstruction (top row), from which we can compute the reconstructed displacement field (middle row) and the corresponding displacement errors (bottom row).

We generate entirely random traction fields via a method the networks have not been trained on. This will prove that our networks indeed have learned to exploit the linearity of the problem and that they have learned a general solution for the problem.

We generate these random traction fields such that they resemble traction fields that my be encountered in reality.

We first generate Gaussian noise with vanishing mean and a standard deviation of $10^{-2}$. We then convolve the substrate field with a proximity filter $\exp(-|\vec{r}-\vec{r}'|^4/0.1^4)$ corresponding to a characteristic correlation length of 0.1 in dimensionless units of the image size, which induces correlations over $\sim 10$ pixels for $N = 104$. The result of this is a traction field and an associated displacement field which can be seen in Fig. S3 (first column). The traction field we have generated here has an entirely different character than the circular training samples the network has seen in training. The displacement field generated from the random traction field is passed to our high noise network and the BFTTC method without further altering it (e.g. by adding noise). The reconstructed traction fields for both methods are shown in Fig. S3 (bottom row). We then pass these reconstructed traction traction fields back into our traction solver, which generates the associated displacement field. The resulting reconstructed displacement fields are shown in Fig. S3 (top row).

Some observations are possible from the visual comparisons alone. First, our network was able to reconstruct the traction field accurately and the resulting reconstructed displacement field is visually similar to the ground truth. Second, the BFTTC method achieves a reconstruction which at first glance seems similar, but on closer inspection clearly shows stronger deviations from the ground truth than our network. Third, the BFTTC method suffers from strong reconstruction artifacts at the perimeter of the substrate. These visual findings are supported by the RMSE we calculate between the ground truth displacement field and the reconstructed displacement fields. While the network has an RMSE $\sim 6.5 \cdot 10^{-2}$ the BFTTC method achieves an RMSE $\sim 1.8 \cdot 10^{-1}$. We generate 50 random force fields and determine the RMSE for the BFTTC method and our $NN_{\text{high}}$ network between the input displacement field and the reconstructed displacement field (generated from the reconstructed traction field). The BFTTC method achieves on average RMSE $\sim 0.04$, while our network achieves on average RMSE $\sim 0.01$. Our network thus predicts traction fields that are more consistent with the input displacement fields.

The advantage of this comparison is that we have the exact ground truth for tractions while not using traction fields that are similar to the training data. We have thus shown two things here – our network is able to reconstruct arbitrary traction fields and it does so with high precision.

### GAUSSIAN NOISE WITH AMPLITUDE CORRELATED WITH POSITION

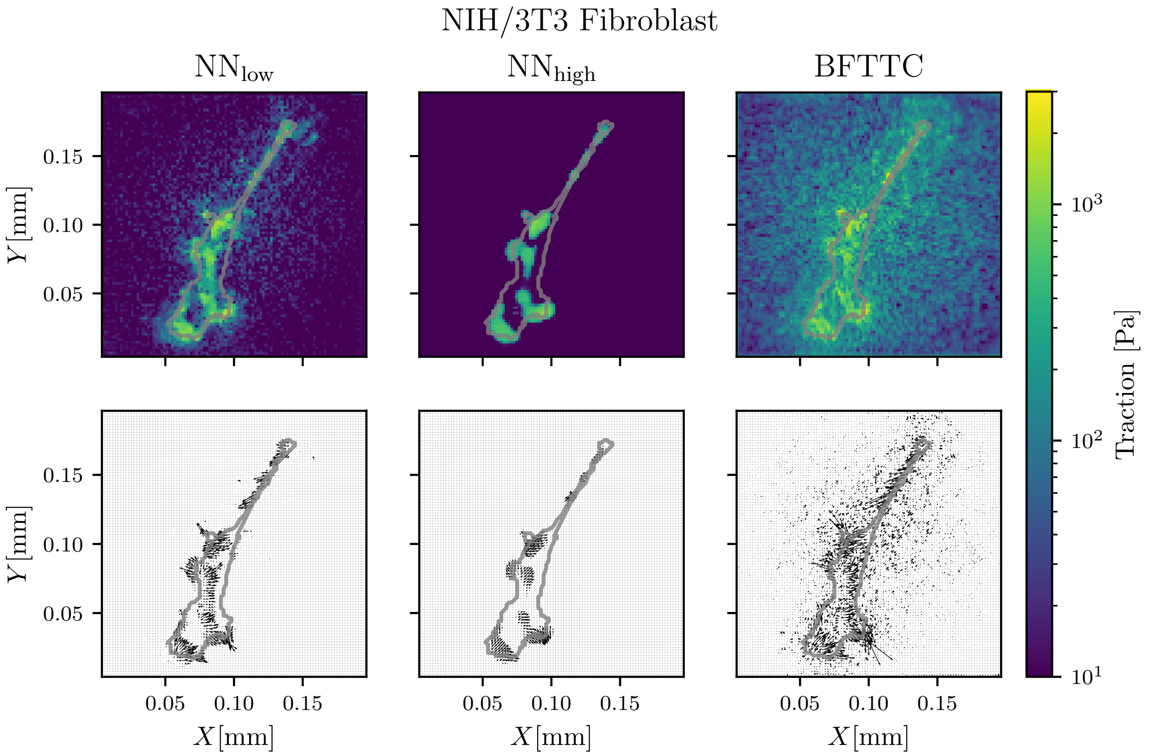


FIG. S4. Results for Gaussian noise with a standard deviation decreasing with the distance to the center of the experimental images. It is evident, that the high noise network returns an invariant prediction even under these circumstances. The BFTTC method produces a significantly elevated traction background noise.

In this work, we focused on uncorrelated Gaussian noise. Experimental noise might contain correlations, for example, from optical aberration that gives rise to a non-uniform noise amplitude. As an example, we consider effects from uncorrelated Gaussian noise with a standard deviation that decreases with the distance from the center of the experimental image in Fig. S4. In this example, the standard deviation decreases with a factor $(1 - r/\sqrt{2})^4$, where $r$ is the dimensionless distance from the center.

### HIGHER TRACTION MAGNITUDES

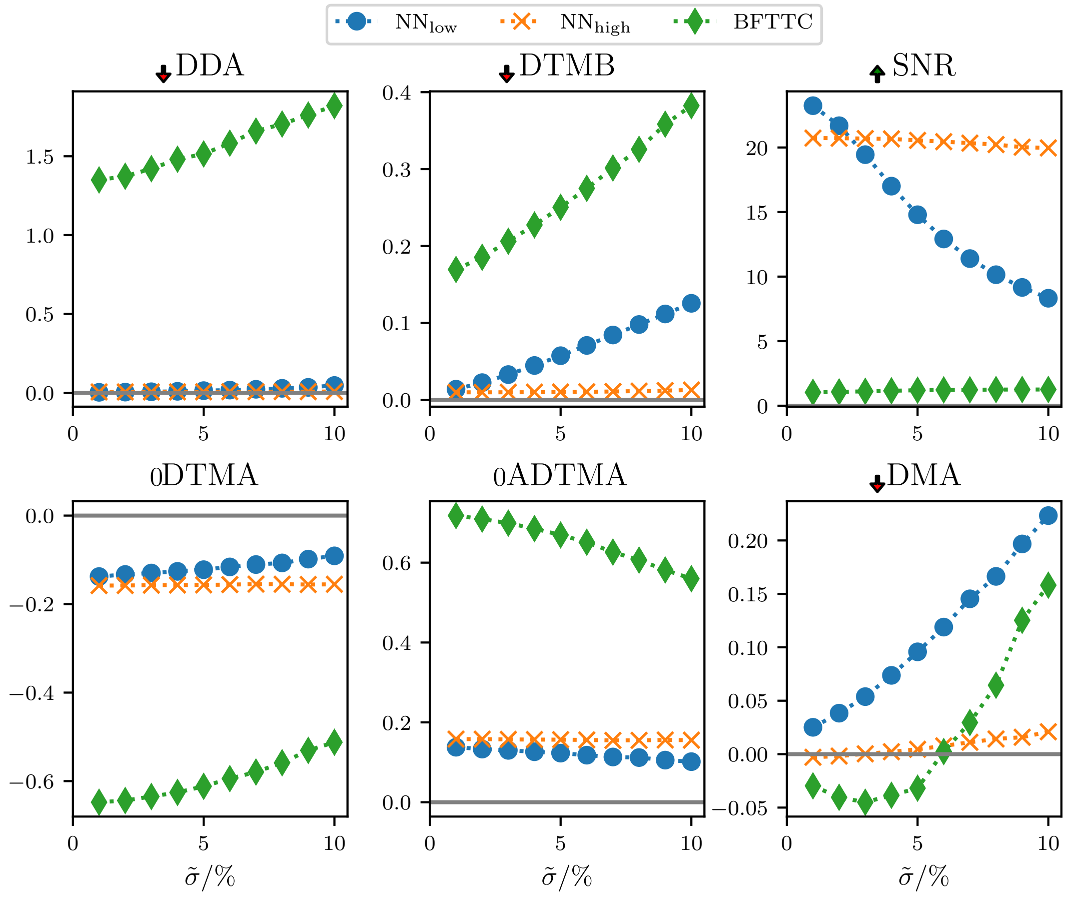


FIG. S5. We generate traction patches with magnitudes uniformly varying from 0.5 to 1.5, which is well outside our training range. We observe that the networks are still able to provide highly accurate predictions, even in this traction magnitude regime.

We trained our networks on traction patches with dimensionless traction magnitudes $\tilde{t}$ uniformly distributed between 0.0 and 0.5. In Fig. S5, we show that the networks are able to generalize to larger traction magnitudes by evaluating all six metrics for synthetic patch-based data with dimensionless traction magnitudes $\tilde{t}$ uniformly distributed between 0.5 and 1.5.

### CHANGING THE INPUT RESOLUTION

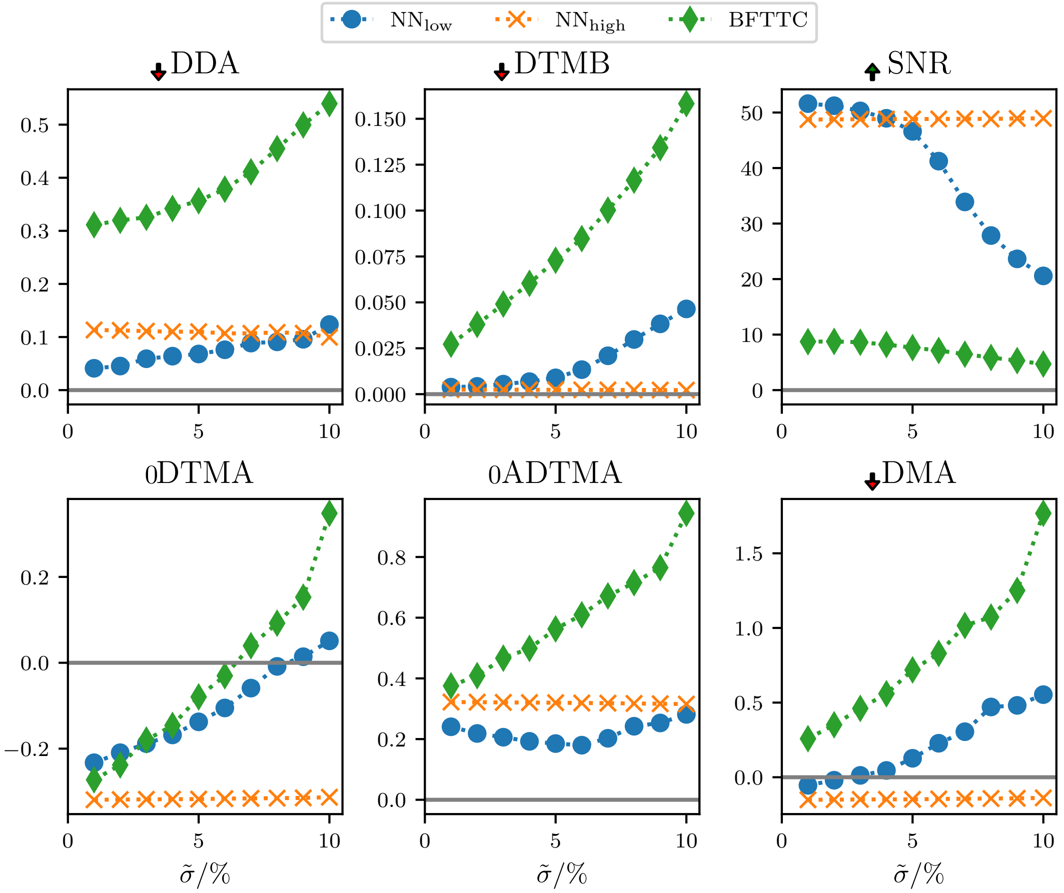


FIG. S6. We generate traction patches on a larger grid of size 256 × 256 to verify that our networks are able to adapt to input resolution changes. We compare our six precision metrics with the BFTTC method and find similar results compared to the findings for 104 × 104 grids in the main text.

To quantify the performance of our networks for higher input resolutions we generate data on a significantly larger grid ($N \times N = 256 \times 256$) and scale the input displacements by 256/104 (as discussed in the main text). We compare the traction reconstruction performance with the BFTTC method in Fig. S6. In our implementation (using TensorFlow), the execution time scales $\propto N^2$ for the BFTTC method, while it remains linear $\propto N$ for network inference.

### FURTHER ANALYSIS OF EXPERIMENTAL DATA

We present additional results for all 14 cells provided in Ref. [S3]. We compare the traction reconstruction for all 14 real Fibroblast between our networks $NN_{low}$ and $NN_{high}$ and the BFTTC method (top and middle rows of each image). Although we do not have the "true" traction field at hand for a quantitative evaluation of precision across the methods, we see compatible results, while both networks have a significantly reduced noise level. The top rows show traction magnitude reconstruction, while the center rows show angle reconstruction. In addition, the bottom rows displays the displacement field computed from the reconstructed traction fields. We see that deviations in the displacement field between all three methods are small. The cells are numbered as in the repository for Ref. [S3]: cell 14 is the cell used as an example in the main text, cells 1-13 are analyzed in addition in this Supporting Information.

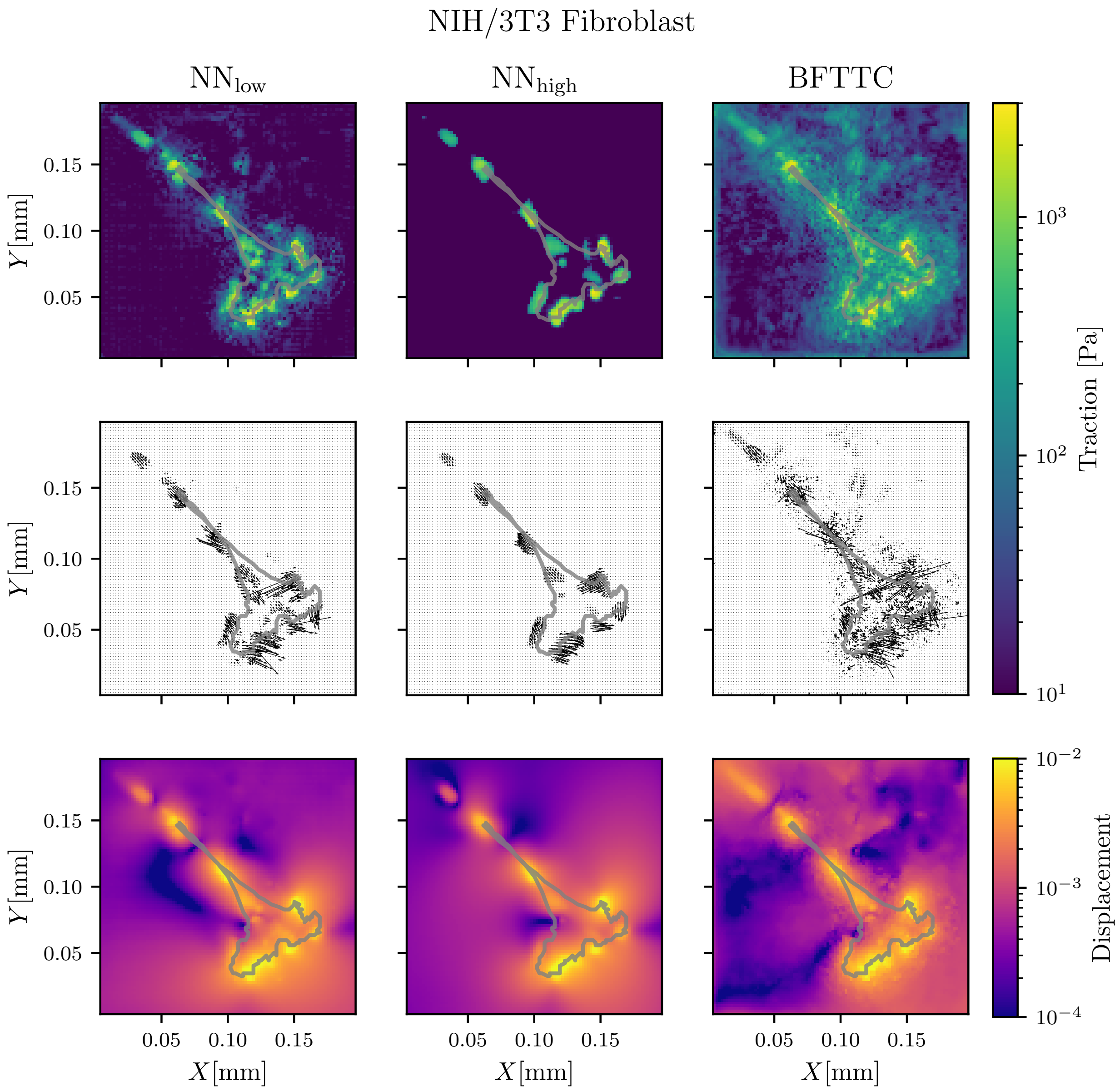


FIG. S7. Comparison of traction and displacement reconstruction for a real Fibroblast between our networks $NN_{low}$ and $NN_{high}$ and the BFTTC method for cell 1 of Ref. [S3].

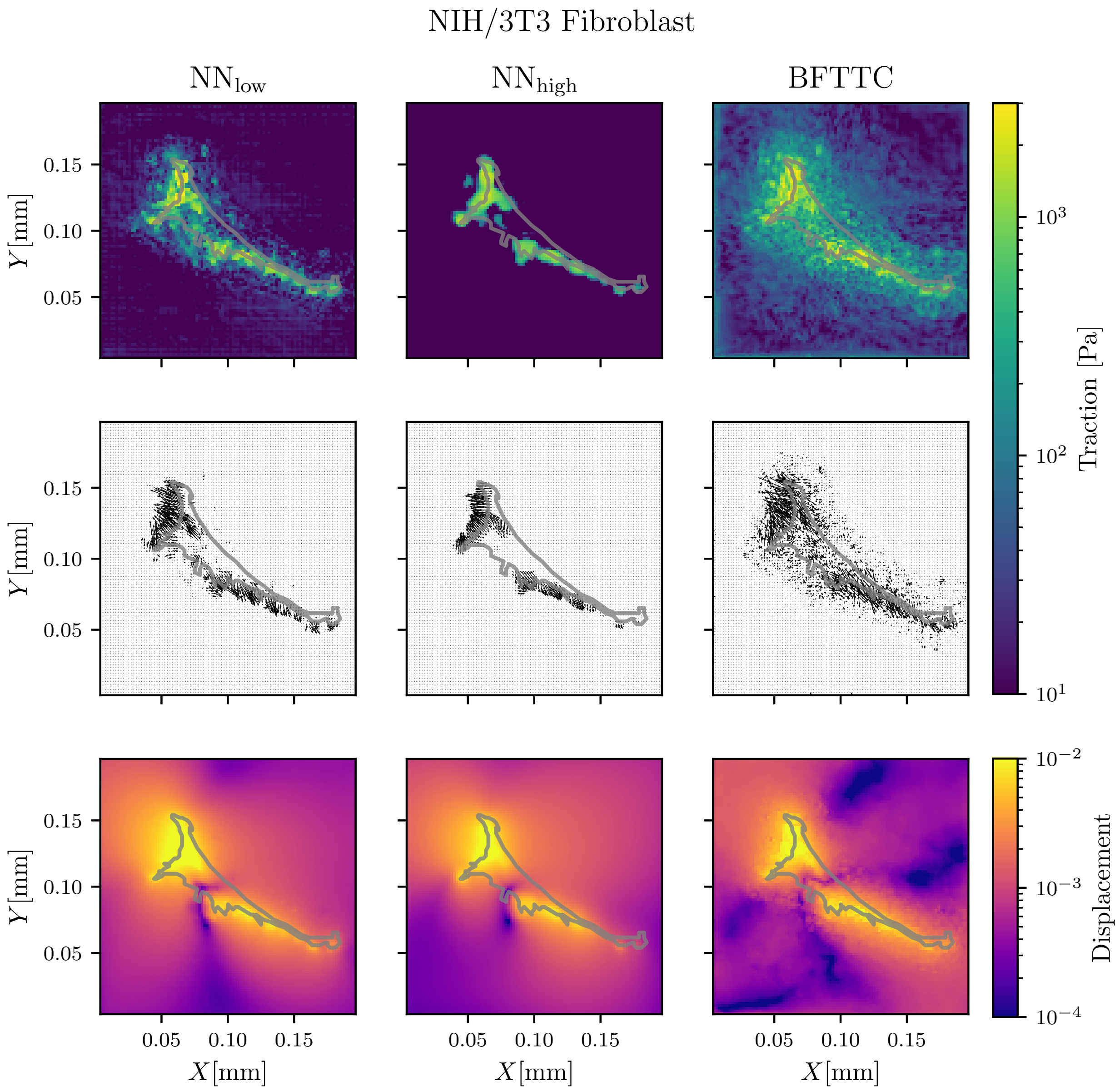


FIG. S8. Comparison of traction and displacement reconstruction for a real Fibroblast between our networks $NN_{low}$ and $NN_{high}$ and the BFTTC method for cell 2 of Ref. [S3].

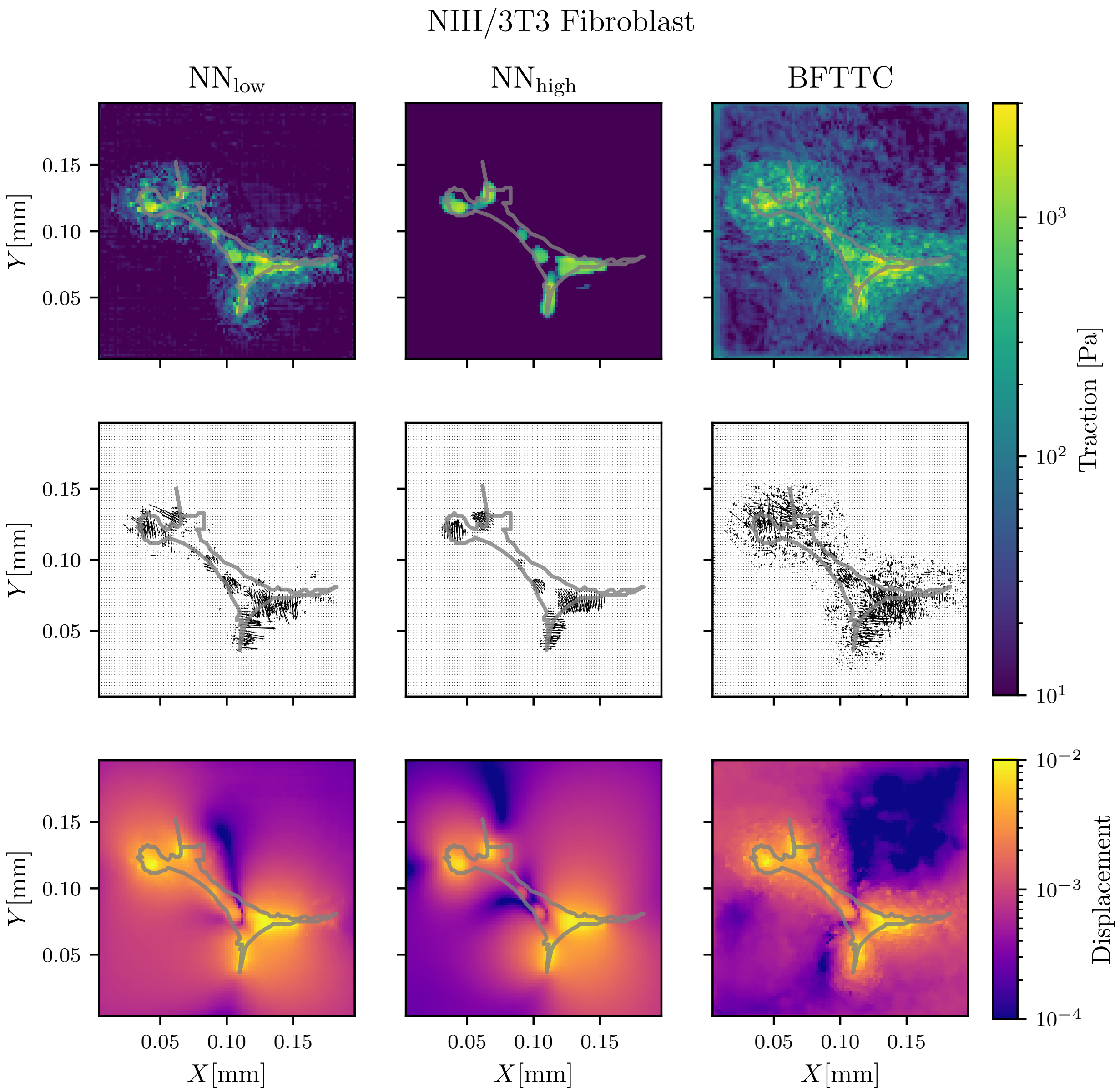


FIG. S9. Comparison of traction and displacement reconstruction for a real Fibroblast between our networks NN$_{low}$ and NN$_{high}$ and the BFTTC method for cell 3 of Ref. [S3].

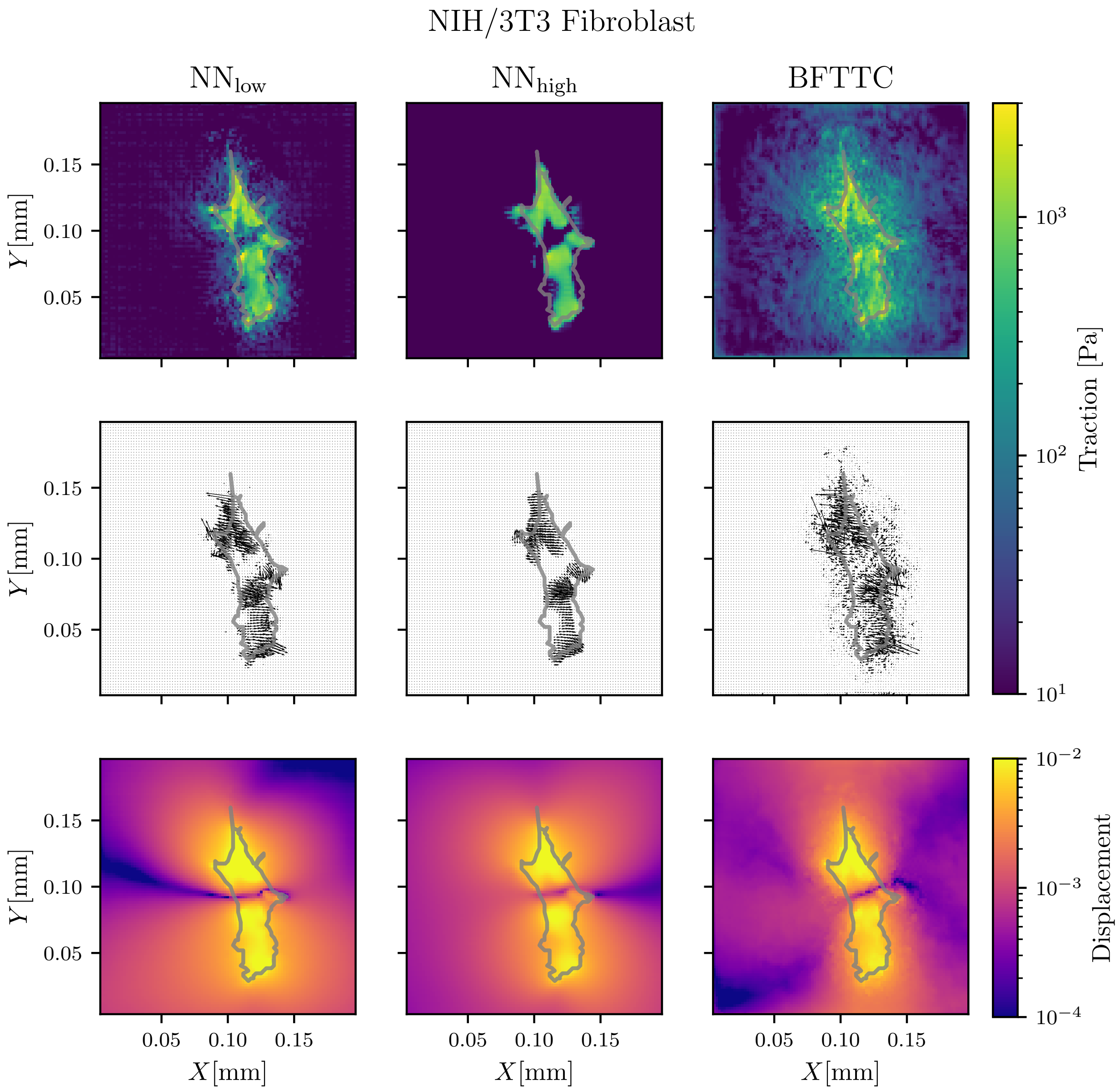


FIG. S10. Comparison of traction and displacement reconstruction for a real Fibroblast between our networks $NN_{low}$ and $NN_{high}$ and the BFTTC method for cell 4 of Ref. [S3].

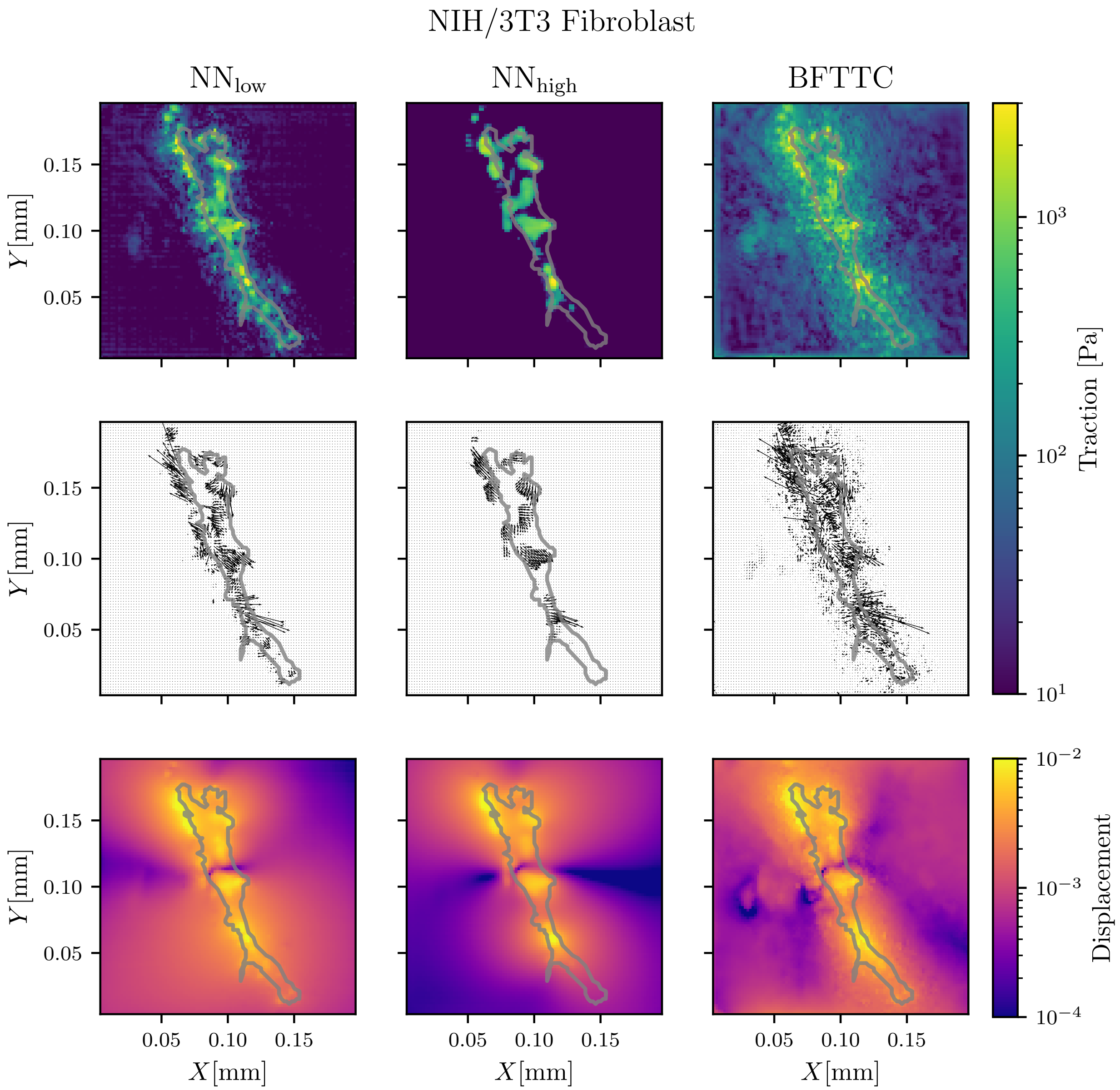


FIG. S11. Comparison of traction and displacement reconstruction for a real Fibroblast between our networks $NN_{low}$ and $NN_{high}$ and the BFTTC method for cell 5 of Ref. [S3].

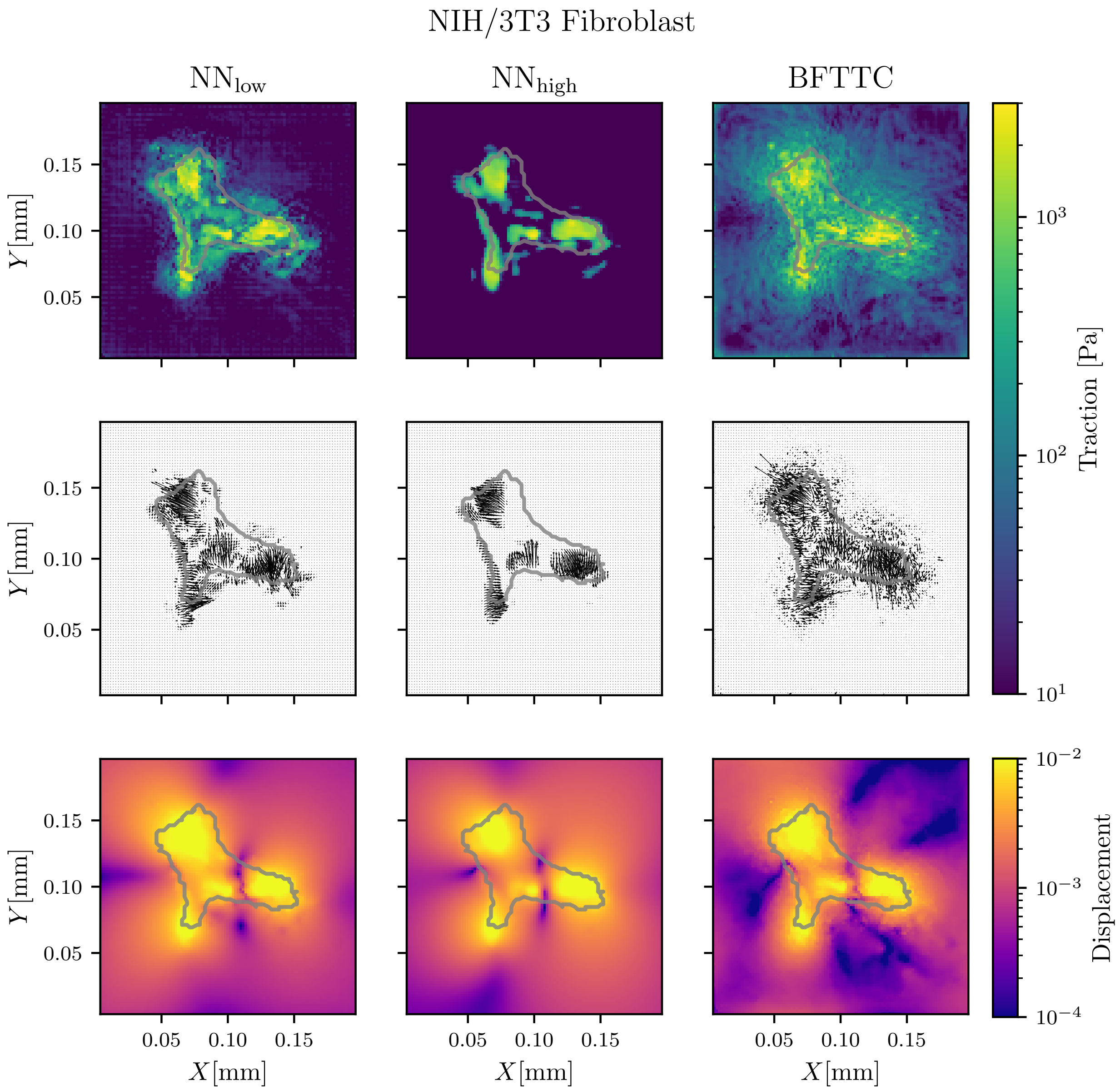


FIG. S12. Comparison of traction and displacement reconstruction for a real Fibroblast between our networks $NN_{low}$ and $NN_{high}$ and the BFTTC method for cell 6 of Ref. [S3].

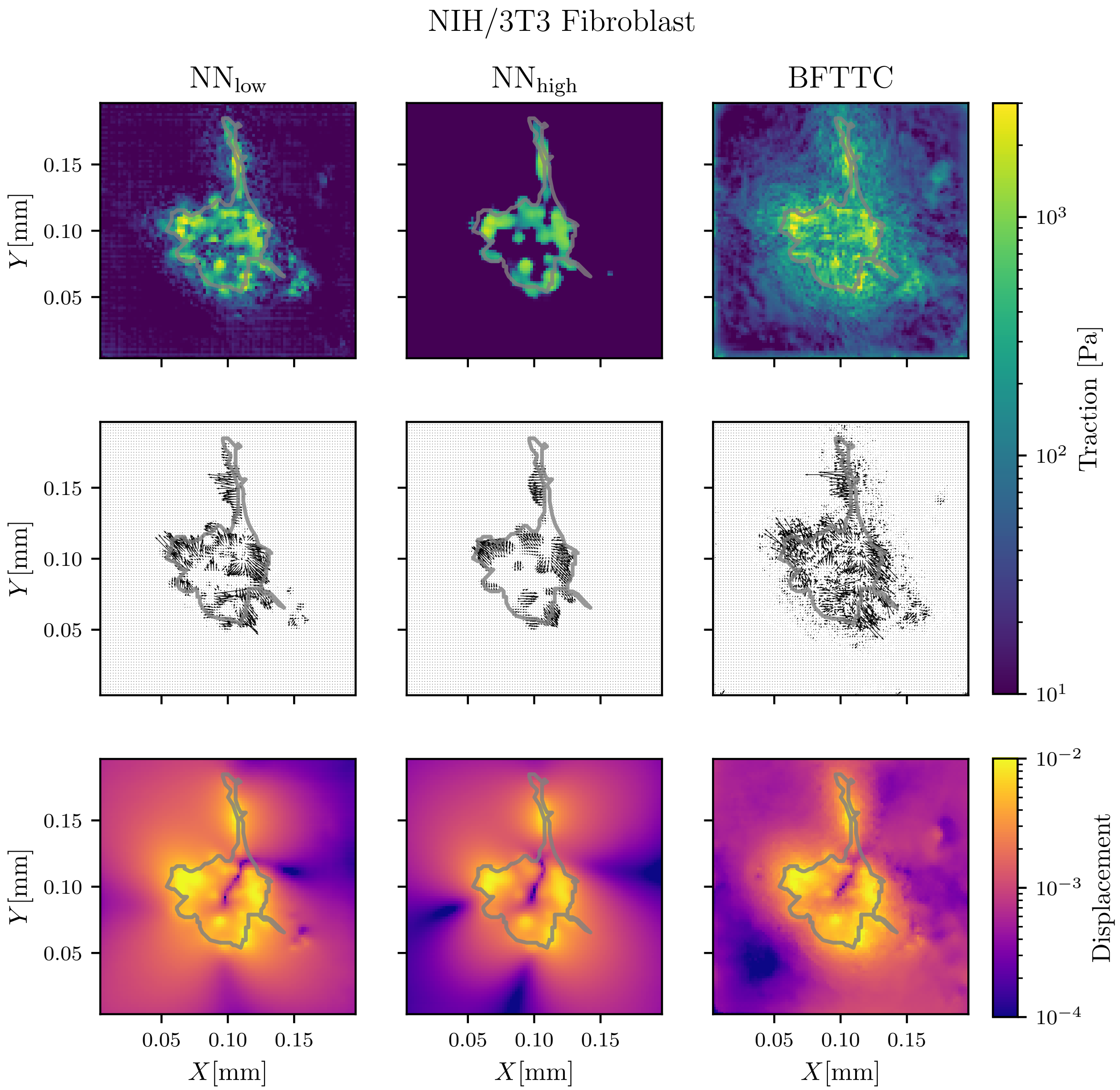


FIG. S13. Comparison of traction and displacement reconstruction for a real Fibroblast between our networks $NN_{low}$ and $NN_{high}$ and the BFTTC method for cell 7 of Ref. [S3].

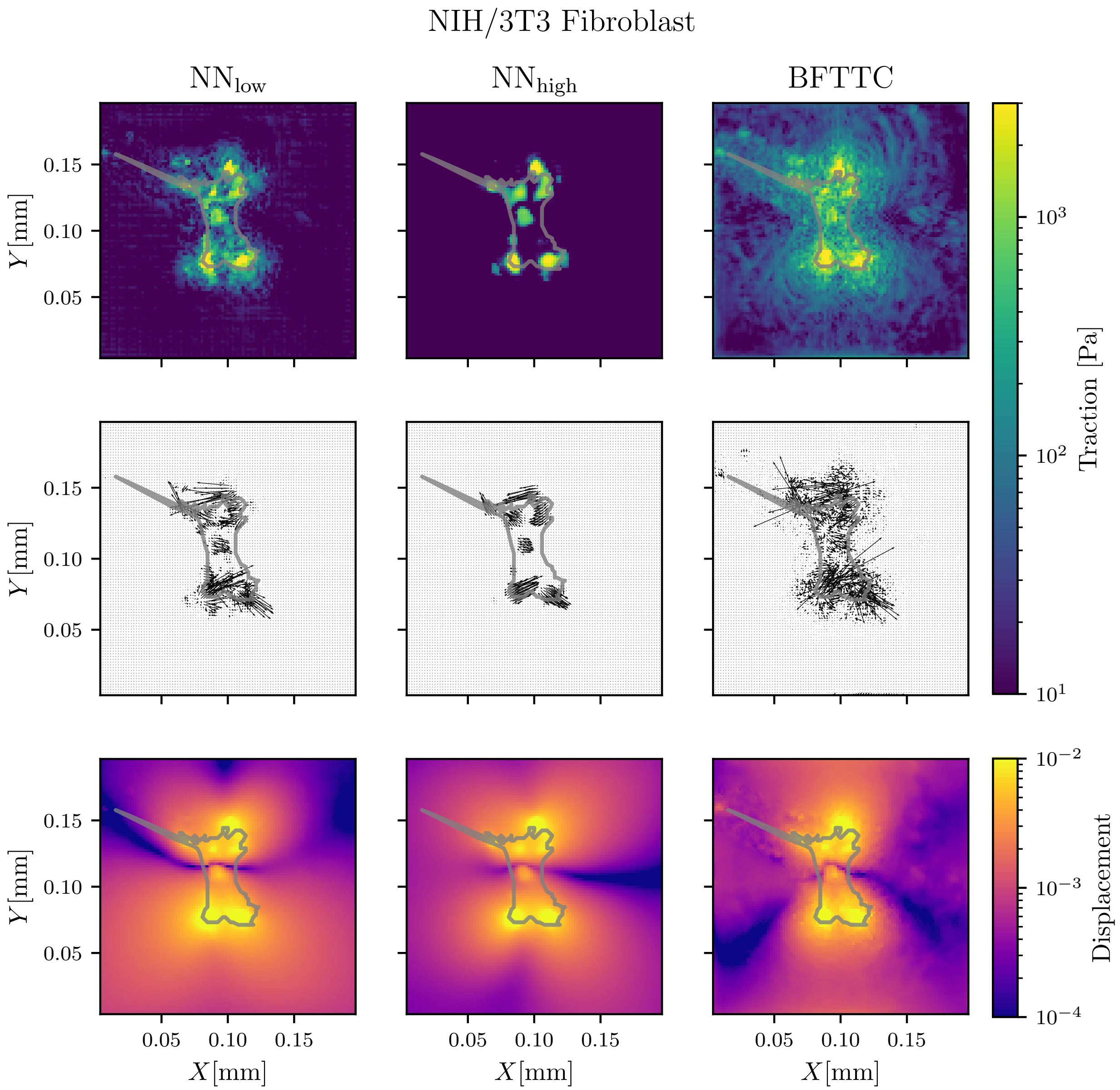


FIG. S14. Comparison of traction and displacement reconstruction for a real Fibroblast between our networks NN$_{low}$ and NN$_{high}$ and the BFTTC method for cell 8 of Ref. [S3].

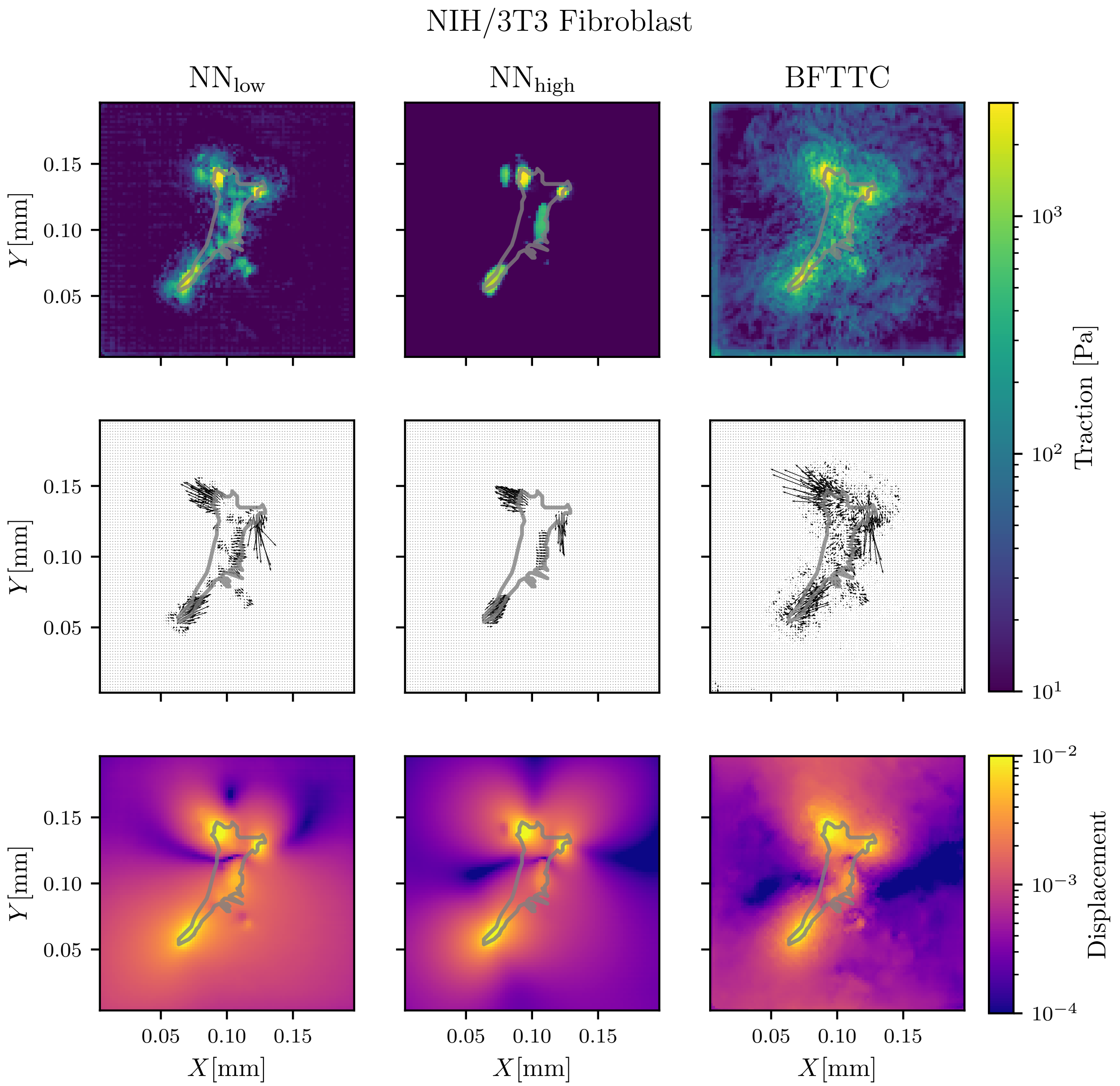


FIG. S15. Comparison of traction and displacement reconstruction for a real Fibroblast between our networks $NN_{low}$ and $NN_{high}$ and the BFTTC method for cell 9 of Ref. [S3].

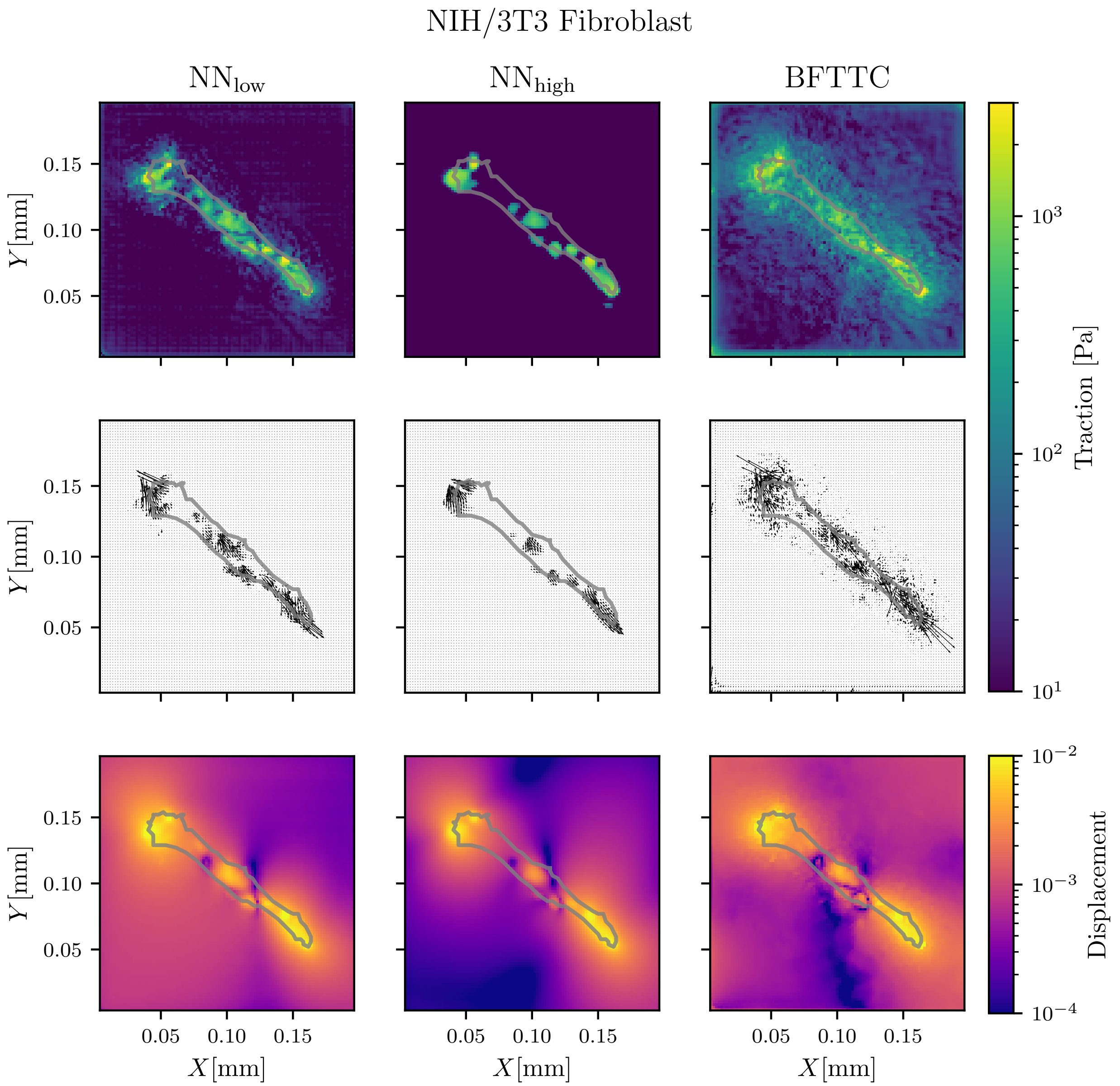


FIG. S16. Comparison of traction and displacement reconstruction for a real Fibroblast between our networks $\mathrm{NN}_{\mathrm{low}}$ and $\mathrm{NN}_{\mathrm{high}}$ and the BFTTC method for cell 10 of Ref. [S3].

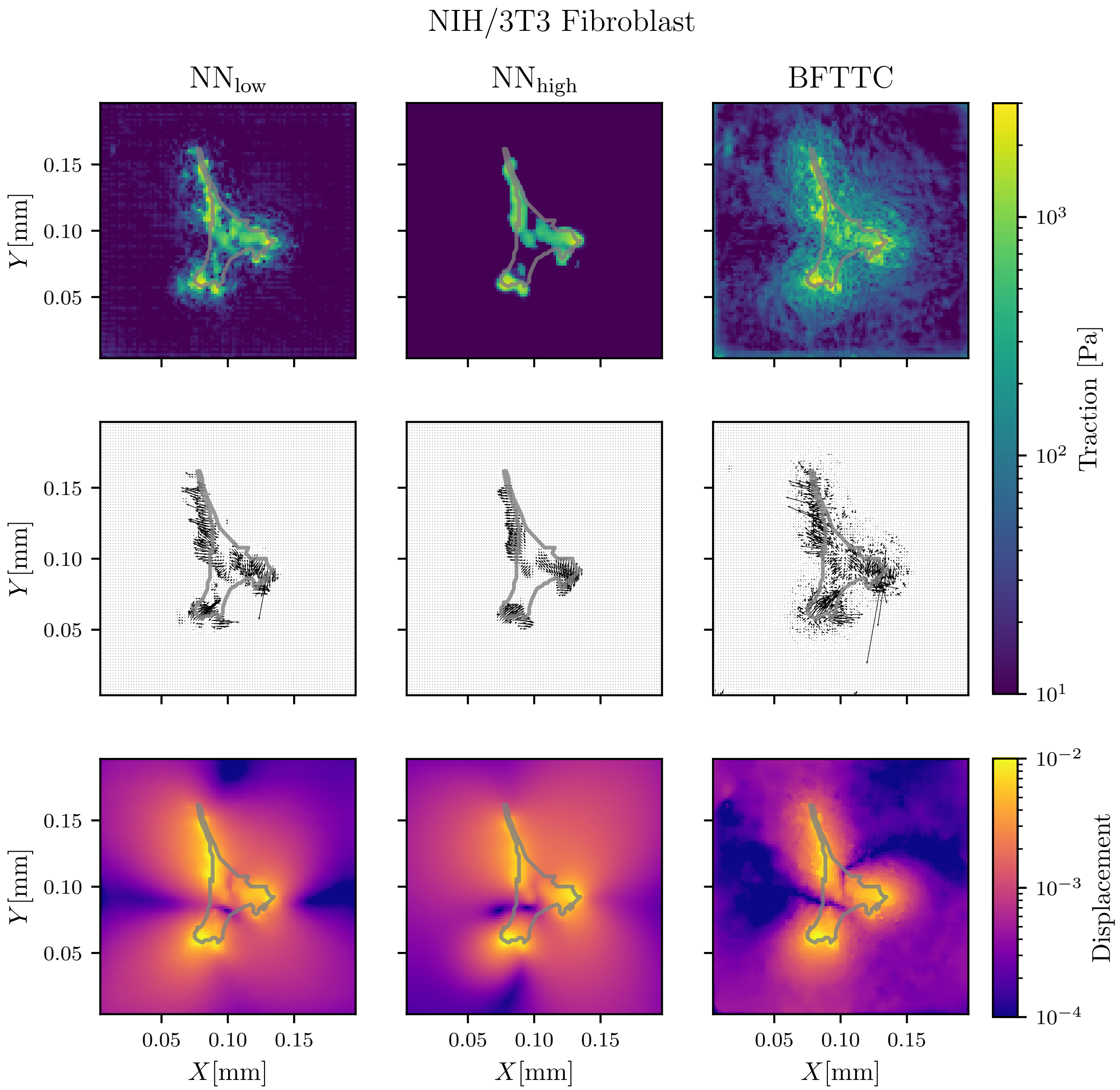


FIG. S17. Comparison of traction and displacement reconstruction for a real Fibroblast between our networks $NN_{low}$ and $NN_{high}$ and the BFTTC method for cell 11 of Ref. [S3].

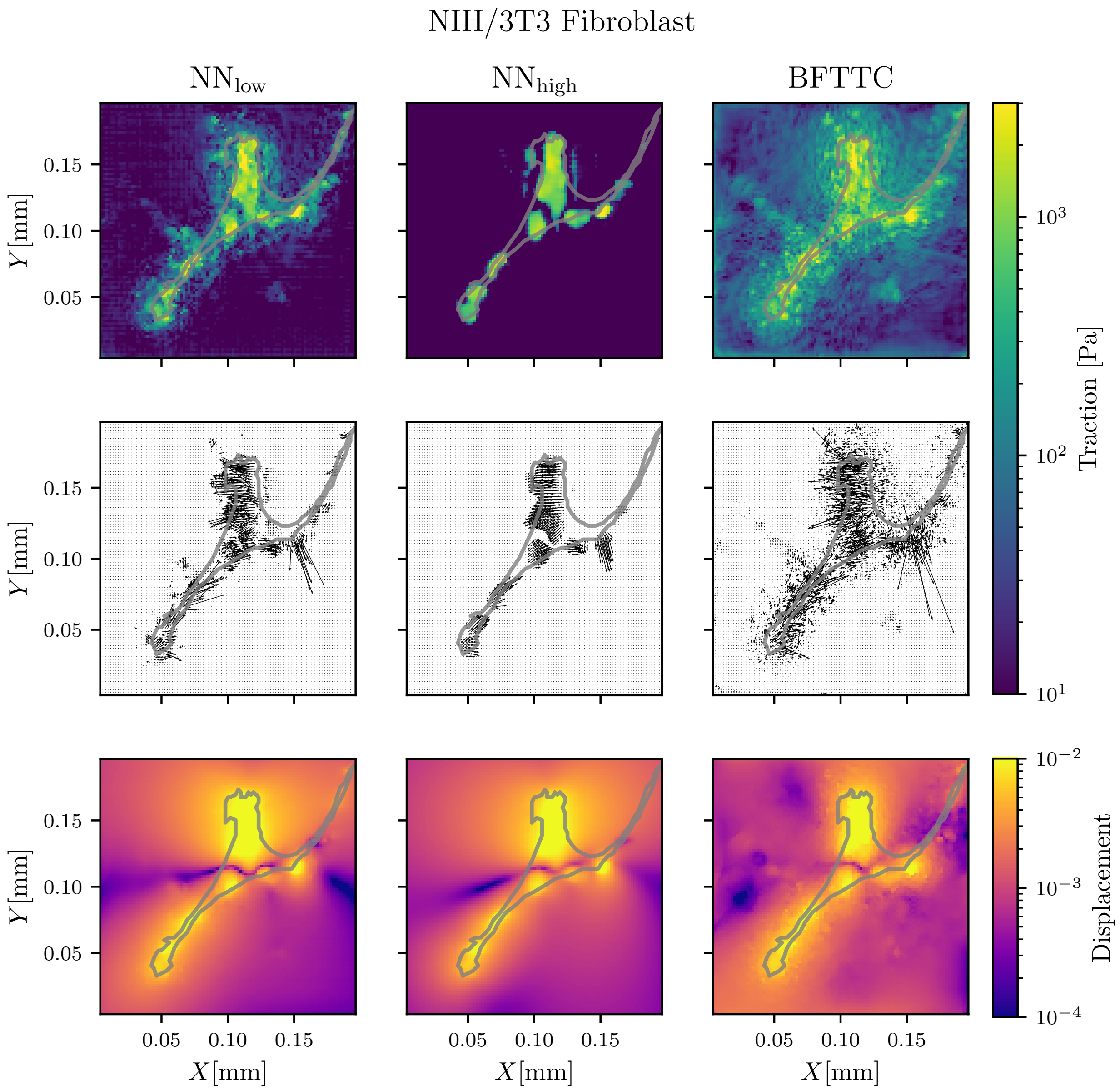


FIG. S18. Comparison of traction and displacement reconstruction for a real Fibroblast between our networks NN$_{\mathrm{low}}$ and NN$_{\mathrm{high}}$ and the BFTTC method for cell 12 of Ref. [S3].

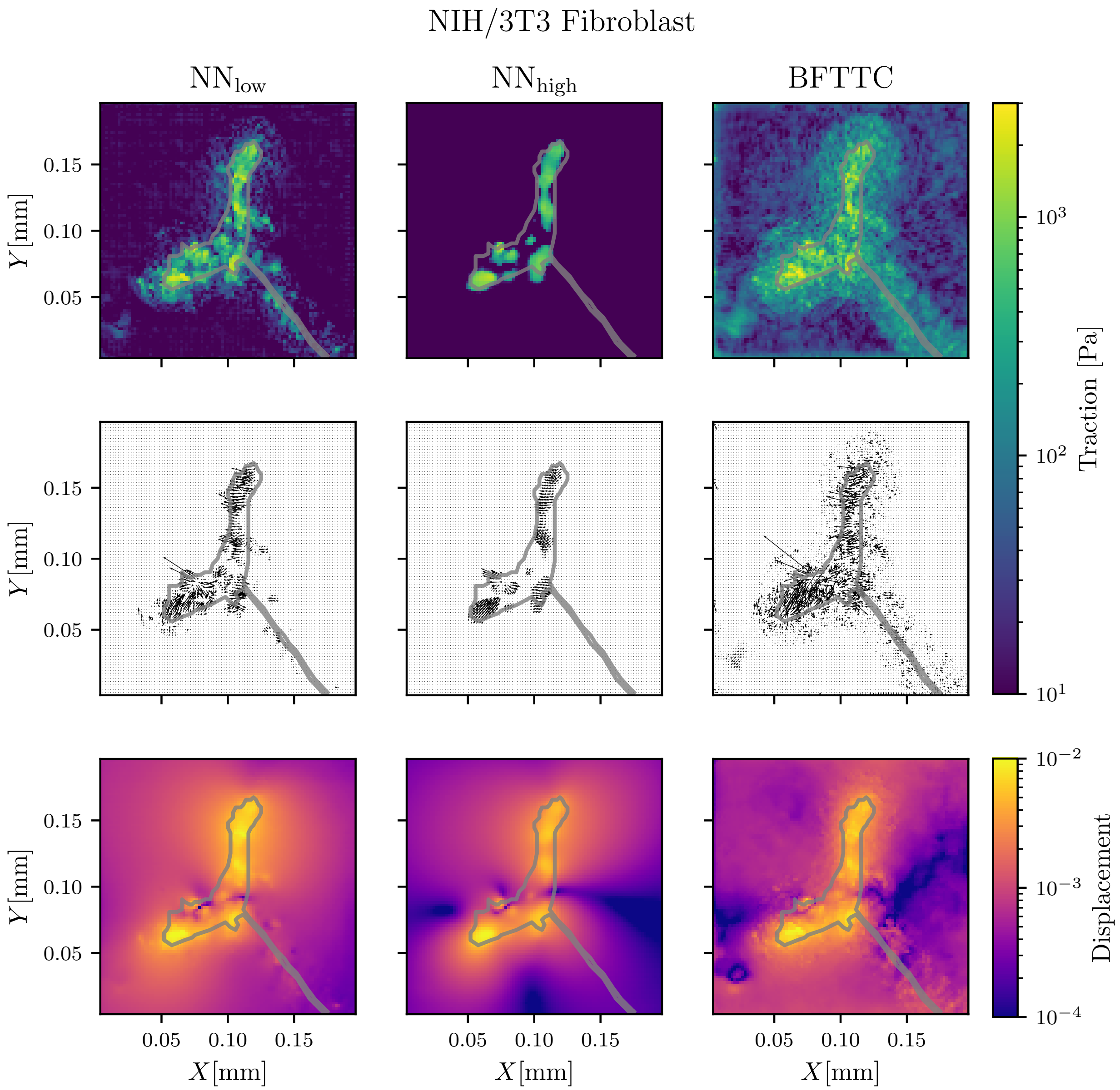


FIG. S19. Comparison of traction and displacement reconstruction for a real Fibroblast between our networks NN$_{\text{low}}$ and NN$_{\text{high}}$ and the BFTTC method for cell 13 of Ref. [S3].

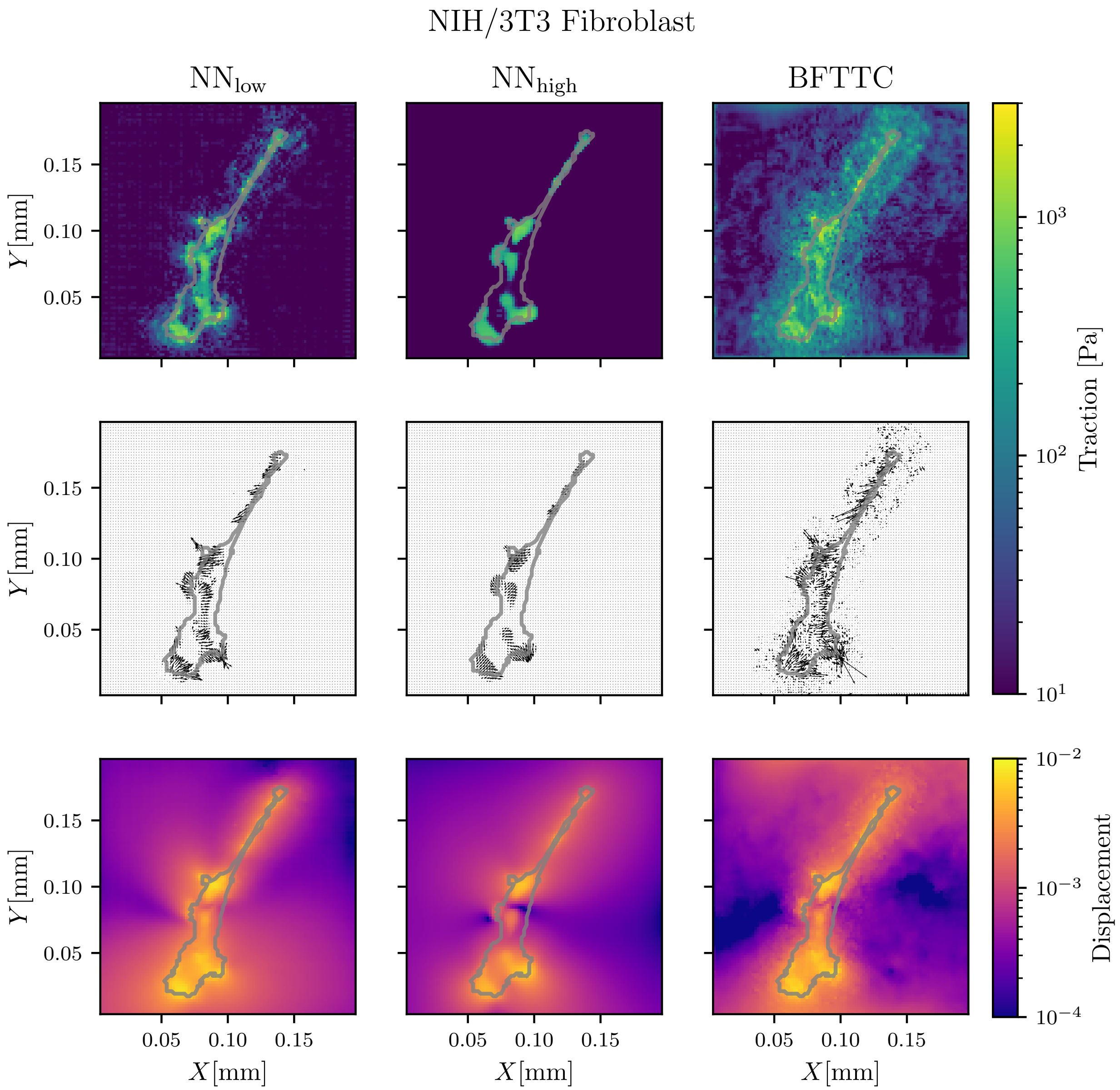


FIG. S20. Comparison of traction and displacement reconstruction for a real Fibroblast between our networks $NN_{low}$ and $NN_{high}$ and the BFTTC method for cell 14 of Ref. [S3]. Cell 14 is also analyzed in Fig. 8 in the main text; here we also show the reconstructed displacement data.

### SUPPORTING REFERENCES

[S1] Landau, L.D. and Lifshitz, E.M. and Kosevitch, A.M. and Pitaevskiĭ, L.P., 1986. Theory of Elasticity. Course of theoretical physics. Pergamon Press, New York.

[S2] Dembo, M., and Y. L. Wang, 1999. Stresses at the cell-to-substrate interface during locomotion of fibroblasts. *Biophys. J.* 76:2307–2316.

[S3] Wang, Y.-l., and Y.-C. Lin, 2021. Traction force microscopy by deep learning. *Biophys. J.* 120:3079–3090.
